# Genomic context sensitizes regulatory elements to genetic disruption

**DOI:** 10.1101/2023.07.02.547201

**Authors:** Raquel Ordoñez, Weimin Zhang, Gwen Ellis, Yinan Zhu, Hannah J. Ashe, André M. Ribeiro-dos-Santos, Ran Brosh, Emily Huang, Megan S. Hogan, Jef D. Boeke, Matthew T. Maurano

## Abstract

Enhancer function is frequently investigated piecemeal using truncated reporter assays or single deletion analysis. Thus it remains unclear to what extent enhancer function at native loci relies on surrounding genomic context. Using the Big-IN technology for targeted integration of large DNAs, we analyzed the regulatory architecture of the murine *Igf2*/*H19* locus, a paradigmatic model of enhancer selectivity. We assembled payloads containing a 157-kb functional *Igf2*/*H19* locus and engineered mutations to genetically direct CTCF occupancy at the imprinting control region (ICR) that switches the target gene of the *H19* enhancer cluster. Contrasting activity of payloads delivered at the endogenous *Igf2*/*H19* locus or ectopically at *Hprt* revealed that the *Igf2*/*H19* locus includes additional, previously unknown long-range regulatory elements. Exchanging components of the *Igf2*/*H19* locus with the well-studied *Sox2* locus showed that the *H19* enhancer cluster functioned poorly out of context, and required its native surroundings to activate *Sox2* expression. Conversely, the *Sox2* locus control region (LCR) could activate both *Igf2* and *H19* outside its native context, but its activity was only partially modulated by CTCF occupancy at the ICR. Analysis of regulatory DNA actuation across different cell types revealed that, while the *H19* enhancers are tightly coordinated within their native locus, the *Sox2* LCR acts more independently. We show that these enhancer clusters typify broader classes of loci genome-wide. Our results show that unexpected dependencies may influence even the most studied functional elements, and our synthetic regulatory genomics approach permits large-scale manipulation of complete loci to investigate the relationship between locus architecture and function.

**HIGHLIGHTS:**

- Composite enhancer elements are subject to genomic context effects mapped to a specific architecture of their endogenous loci.
- *Igf2/H19* expression is affected by long-range regulatory elements beyond the canonically defined locus, and the *H19* enhancer cluster in particular relies on the surrounding context at its endogenous locus.
- The *Sox2* LCR functions as an autonomous enhancer without requiring additional surrounding context.
- The influence of genomic context is buffered at intact loci, but manifests more strongly as key regulatory elements are deleted or repositioned.

## INTRODUCTION

Genomic regulatory elements are often treated as discrete peaks, but their collective activity establishes distinct genomic neighborhoods with widely varying regulatory element density and position relative to genes. Cross-tissue DNase I hypersensitive site (DHS) patterns frequently resemble the activity of their target genes (Thurman et al. 2012), suggesting a role for genomic context in DHS function. Since distinct expression programs often co-occupy nearby genomic space, genomic regulatory architecture must provide for a degree of selectivity. Selective expression of certain genes might be facilitated through promoter-enhancer compatibility (Bergman et al. 2022; Martinez-Ara et al. 2022), competition for regulatory elements, interposing regulatory elements or genomic distance (Peterson and Stamatoyannopoulos 1993; Zuin et al. 2022), or boundary elements that confer compartmentalization and large-scale domain organization (Symmons et al. 2014; Ringel et al. 2022). Questions of context and sufficiency have been studied in a genomic context at the level of individual DHSs or transcription factor (TF) recognition sequences (Bolt et al. 2022; Blayney et al. 2023; Brosh et al. 2023). Yet enhancer elements are typically investigated through limited genome engineering at their endogenous loci, and the lack of methods for genome editing of regions upwards of 10 kb has resulted in a limited understanding of locus architecture and genomic context at larger length scales.

Studies of simpler reporter assays have shown that expression is modulated by genomic context composed of quantitative contributions of many elements acting at ranges upwards of 100 kb (Maricque et al. 2019; Hong and Cohen 2022; Ribeiro-Dos-Santos et al. 2022). Certain composite regulatory elements such as locus control regions (LCR) confer position-independent expression, and super-enhancers are argued to have properties beyond those of their constituent enhancers (Li et al. 2002; Moorthy et al. 2017; Blobel et al. 2021). Autonomy might result from inclusion of multiple regulatory elements to provide resilience, specific genomic features such as insulation or domains, or simply additional intervening distance from other surrounding regulatory elements. Thus it is possible that complex regulatory elements have evolved to permit more precise control over these features. However, the exact sequence elements required to confer robust locus activity remain unclear.

The canonical *Igf2*/*H19* imprinted locus includes multiple complex regulatory elements and is a model for enhancer selectivity (Arney 2003). The insulin-like growth factor 2 (*Igf2*) gene is expressed from the paternal allele while the non-coding *H19* gene is maternally expressed. The “boundary model” posits that an imprinting control region (ICR) located between the two genes directs the activity of a shared enhancer cluster (**Figure 1A**) (Bell and Felsenfeld 2000; Hark et al. 2000). This ICR contains a tandem array of CTCF sites which are methylated and unoccupied by CTCF on the paternal allele, allowing the enhancers to interact with and activate the more distal *Igf2* gene. The maternal allele is unmethylated and occupied by CTCF, which acts as an insulator that directs the enhancer cluster to activate the proximal *H19* instead of *Igf2* (**Figure 1B**). A transgenic YAC containing a 130-kb sequence from the *Igf2*/*H19* locus has been shown sufficient to recapitulate this pattern of imprinted gene expression when integrated randomly (Ainscough et al. 1997). Therefore, the *Igf2*/*H19* bimodal regulation offers a unique paradigm to understand how locus architecture and genomic context might influence the activity of the individual regulatory components in a specific locus and their interactions with their regulatory landscape.

**Figure 1.**
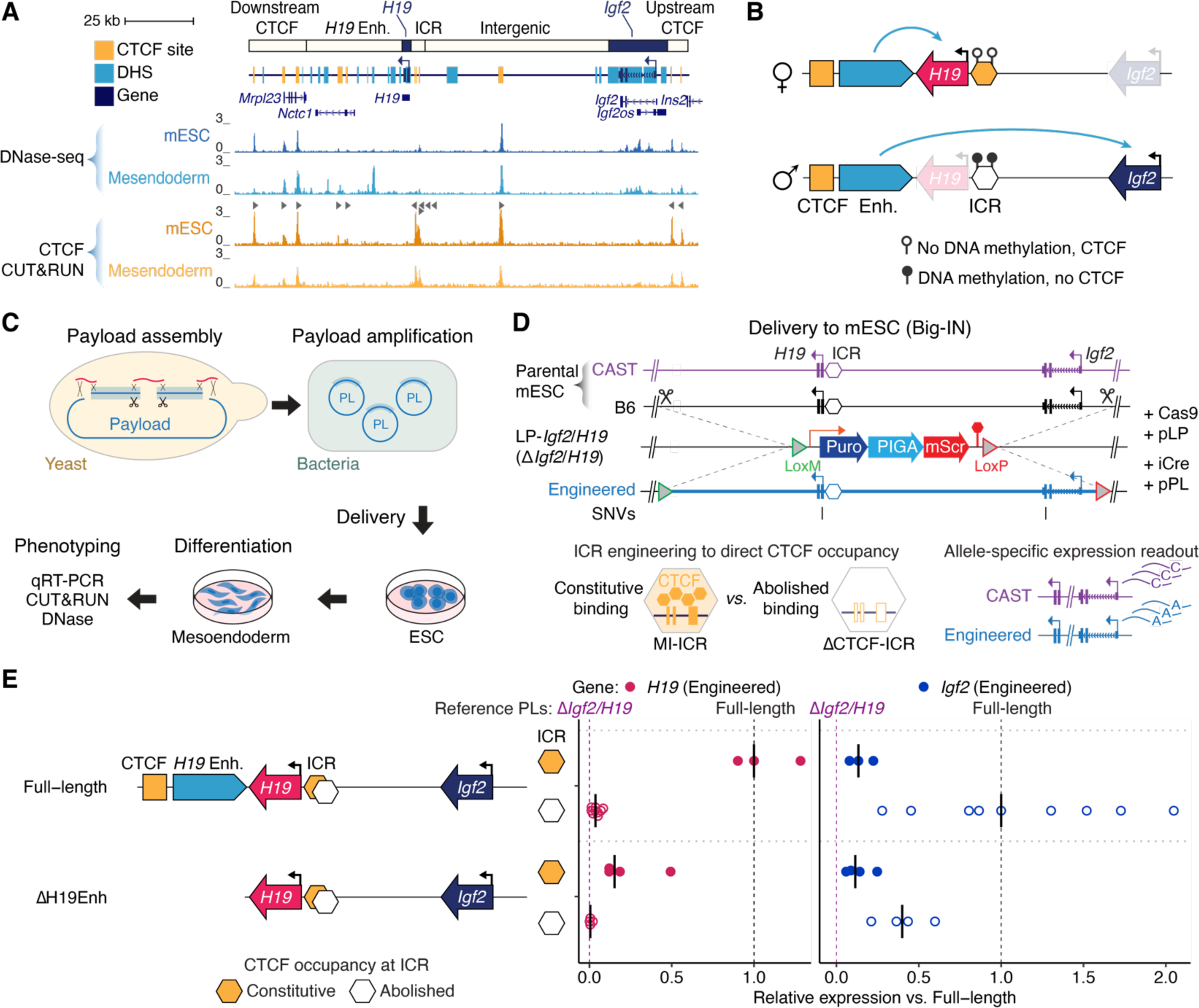
Engineering the *Igf2/H19* locus. (**A**) *Igf2/H19* locus in mESC and differentiated mesendodermal cells. DNase-seq tracks show activation of the enhancer cluster upon differentiation. Triangles indicate CTCF recognition sequence orientation. (**B**) Barrier model where CTCF occupancy at the imprinting control region (ICR) directs activity of the endogenous enhancer cluster towards *H19* or *Igf2* (Bell and Felsenfeld 2000; Hark et al. 2000). (**C**) Schematic of engineering and delivery strategy. Payloads were assembled in yeast, amplified in bacteria, and delivered to mESCs using Big-IN. Multiple independent clones were systematically verified for correct genome engineering and differentiated into mesendodermal cells for further phenotyping. (**D**) Schematic of *Igf2/H19* locus engineering using Big-IN. CpG mutations (methylation-insensitive ICR [MI-ICR]) or CTCF deletions (ΔCTCF-ICR) permit genetic control of CTCF occupancy. B6 or engineered variants in *H19* and *Igf2* genes permit allele-specific expression analysis. (**E**) Expression analysis of the engineered *Igf2/H19* locus in mesendodermal cells. Schemes on the left represent each delivered payload. At right, each point represents the expression of the engineered *H19* or *Igf2* gene in an independent clone, measured using allele-specific qRT-PCR. Bars indicate median. Expression was normalized to *Gapdh* and then scaled between 0 (Δ*Igf2/H19* locus) and 1 (“Full-length” rescue payloads), normalizing *H19* to the “Full-length” payload with constitutive CTCF occupancy, and *Igf2* to the “Full-length” payload with abolished CTCF occupancy (dashed lines). Payloads with both constitutive CTCF occupancy (orange hexagon) or abolished CTCF occupancy (empty hexagon) at the ICR were assessed.

Here we take advantage of our Big-IN technology (Brosh et al. 2021) for repeated delivery of large DNA sequences to develop a synthetic regulatory genomics approach to quantitatively dissect enhancer cluster context sensitivity. Using this approach to deliver payloads up to 158 kb in length, we identified a new long-range distal enhancer cluster at the *Igf2*/*H19* locus that, together with the endogenous *H19* enhancers, directs *Igf2* and *H19* expression in an ICR-dependent manner. Replacement of the well-described *Sox2* LCR by the *H19* enhancer cluster shows that full function of the latter relies on additional elements at its endogenous locus. In contrast, the *Sox2* LCR can function outside its native context to drive robust expression of both *Igf2* and *H19*. However, the *Sox2* LCR responds only partially to the ICR, as it is able to maintain high *H19* expression levels despite CTCF occupancy. We show that the independence of the *Sox2* LCR and the context dependence of the *H19* enhancers are reflected in cross-cell type patterns of regulatory DNA actuation which manifest broadly at many other loci genome-wide, providing clues to the regulatory architecture of these and other important developmentally regulated genes.

## RESULTS

### A synthetic *Igf2/H19* locus as a model for dissection of enhancer selectivity

To quantitatively dissect the sensitivity of enhancer elements to their broader genomic context, we developed a synthetic regulatory genomics approach to manipulate the *Igf2*/*H19* locus in cultured cells. We employed our Big-IN technology (Brosh et al. 2021) for repeated delivery (**Figure 1C**) to mouse embryonic stem cells (mESC). To facilitate allele-specific engineering and readout, we employed a C57BL/6J (B6, maternal) × CAST/EiJ (CAST, paternal) mESC line derived from an *M. musculus* and *M. musculus castaneus* F1 hybrid mouse, which harbors heterozygous point variants every 140 bp on average throughout its genomic sequence (Keane et al. 2011; Eckersley-Maslin et al. 2014). We first installed a Big-IN landing pad (LP) replacing the *Igf2/H19* locus on the B6 allele (**Figure 1D**). After comprehensive quality control using PCR genotyping and targeted capture sequencing (Capture-seq), we identified clones with single-copy, on-target landing pad integration, named *LP-Igf2/H19* mESCs (**Figure S1**).

We then assembled in yeast a baseline payload containing a complete 157-kb *Igf2/H19* locus into a Big-IN delivery vector, starting from a B6 mouse-derived bacterial artificial chromosome (BAC). From this baseline payload, we derived a set of further engineered derivatives (**Figure S2**, **Table S1**). Payloads were sequence-verified (**Table S2**) and delivered to *LP-Igf2/H19* mESCs using Big-IN (**Figure 1D**). Multiple mESC clones were isolated for each payload and comprehensively verified for single-copy targeted payload integration using PCR genotyping and Capture-seq (**Table S3**).

As both *Igf2* and *H19* are transcriptionally inactive in mESCs, we employed a mesendodermal differentiation protocol (Thomson et al. 2011) which induces robust expression of both *Igf2* and *H19* (**Figure S3**) and increased DNA accessibility at the enhancer cluster located downstream of *H19* (**Figure 1A**). RNA-seq analysis to identify differentially expressed genes between mESCs and differentiated mesendodermal cells confirmed that previously identified (Shen et al. 2008) genes upregulated in mesendoderm were enriched and genes downregulated were depleted (**Table S4**, **Figure S4A-B**). From the RNA-seq data, we identified a panel of 5 genes upregulated in mesendoderm and designed qRT-PCR assays to measure their expression across all samples in this study (**Figure S4C**). Principal component analysis (PCA) of mesendoderm marker gene expression showed that the mESC and differentiated cells were clearly separated (**Figure S4D**).

While stable parent-of-origin specific methylation is established at the *Igf2*/*H19* ICR during development, the methylation pattern maintained in cultured mESCs is more labile (Swanzey et al. 2020). In addition, controlling methylation status of transfected DNA would require additional steps (Schübeler et al. 2000; Butz et al. 2022). Thus we developed a genetic approach to reliably direct CTCF occupancy at the ICR (**Figure 1D**). We engineered a methylation-insensitive ICR (MI-ICR) where CpGs were mutated in all 6 CTCF recognition sequences without reducing predicted CTCF binding energy (**Figure S5A-B**). Similarly, we engineered a ΔCTCF-ICR where all 6 CTCF recognition sequences were surgically deleted (**Figure S5A**). We profiled both parental and engineered mESCs using CUT&RUN to verify the expected CTCF occupancy. In parental cells, we detected biallelic CTCF occupancy throughout the locus, except at the ICR, where only the maternal B6 allele was occupied. Delivery of a wild-type ICR or a MI-ICR payload resulted in specific CTCF occupancy on the engineered allele, while the un-engineered CAST allele remained unoccupied. In contrast, CTCF occupancy was absent in ΔCTCF-ICR payloads, demonstrating that our synthetic ICR sequences can genetically direct CTCF occupancy independently of parental origin of the engineered chromosome (**Figure S5C**).

We developed an allele-specific qRT-PCR strategy for expression analysis of the engineered allele. We designed two point variants in the *H19* transcript sequence to distinguish the engineered and CAST alleles, and four synonymous point variants in the *Igf2* transcript sequence (**Figure 1D**, **Figure S6**). We performed a ΔCt analysis using *Gapdh* as a reference, and scaled *Igf2* and *H19* values between 0 (deletion of the engineered allele, Δ*Igf2/H19*) and 1 (normalizing *H19* to the “Full-length” payload with constitutive CTCF occupancy, and *Igf2* to the “Full-length” payload with abolished CTCF occupancy). This approach allowed us to rewrite the *Igf2*/*H19* locus for functional dissection and permitted a precise transcriptional readout of the engineered allele.

Expression analysis of the “Full-length” payloads demonstrated reciprocal *Igf2* and *H19* expression dependent upon CTCF occupancy at the ICR, with higher *H19* expression in the presence of CTCF, and higher *Igf2* expression in absence of CTCF (**Figure 1E, Figure S7, Table S5**). Thus, our experimental system recapitulates the boundary model of the *Igf2/H19* locus (Bell and Felsenfeld 2000; Hark et al. 2000) (**Figure 1B**). Surprisingly, we observed that the “ΔH19Enh” payloads, while completely lacking the *H19* enhancer cluster, still retained modest expression of both *Igf2* and *H19* (**Figure 1E**), suggesting that the canonical *H19* enhancers are not the only determinant of gene expression at this locus.

To ensure reproducibility of our results, we analyzed several independent clones for each delivered payload to account for potential epigenetic variability during delivery and differentiation. A subset of clones were differentiated multiple times and showed high concordance in expression of both mesendoderm marker genes and engineered alleles (**Figure S8**). We compared the variability of the same payload in multiple independently isolated clones and found it similar to that across multiple differentiations (**Figure S9**). While there have been suggestions of feedback between the two alleles at this locus (Sandhu et al. 2009), *Igf2* expression in our system was not correlated between the engineered and CAST alleles (**Figure S10**).

### Dissection of the *Igf2/H19* regulatory architecture reveals a novel distal enhancer cluster

We thus investigated whether additional regulatory elements outside the engineered region play a role in *Igf2/H19* regulation (**Figure 2A**). Inversion of a locus preserves its internal organization but alters the polarity and distance of interactions with surrounding elements. We thus engineered payloads with an inverted *Igf2*/*H19* locus and delivered them to LP-*Igf2/H19* mESCs (**Figure 2B**). These “Full-length inverted” payloads showed strongly reduced *H19* expression, while “ΔH19Enh inverted” payloads showed completely ablated *H19* expression. These results indicate that *H19* expression is highly sensitive to regulatory elements beyond the payload boundary. In contrast, inversion had little effect on *Igf2* levels in the payloads with abolished CTCF occupancy. In payloads with constitutive CTCF occupancy, *Igf2* expression even increased slightly. As inverted payloads showed increased expression of *Igf2* and reduced expression of *H19*, our results imply the presence of regulatory elements downstream of the canonical 157-kb locus which cooperate with known regulatory elements and exhibit differing influences on *H19* and *Igf2* (**Figure 2A**).

**Figure 2.**
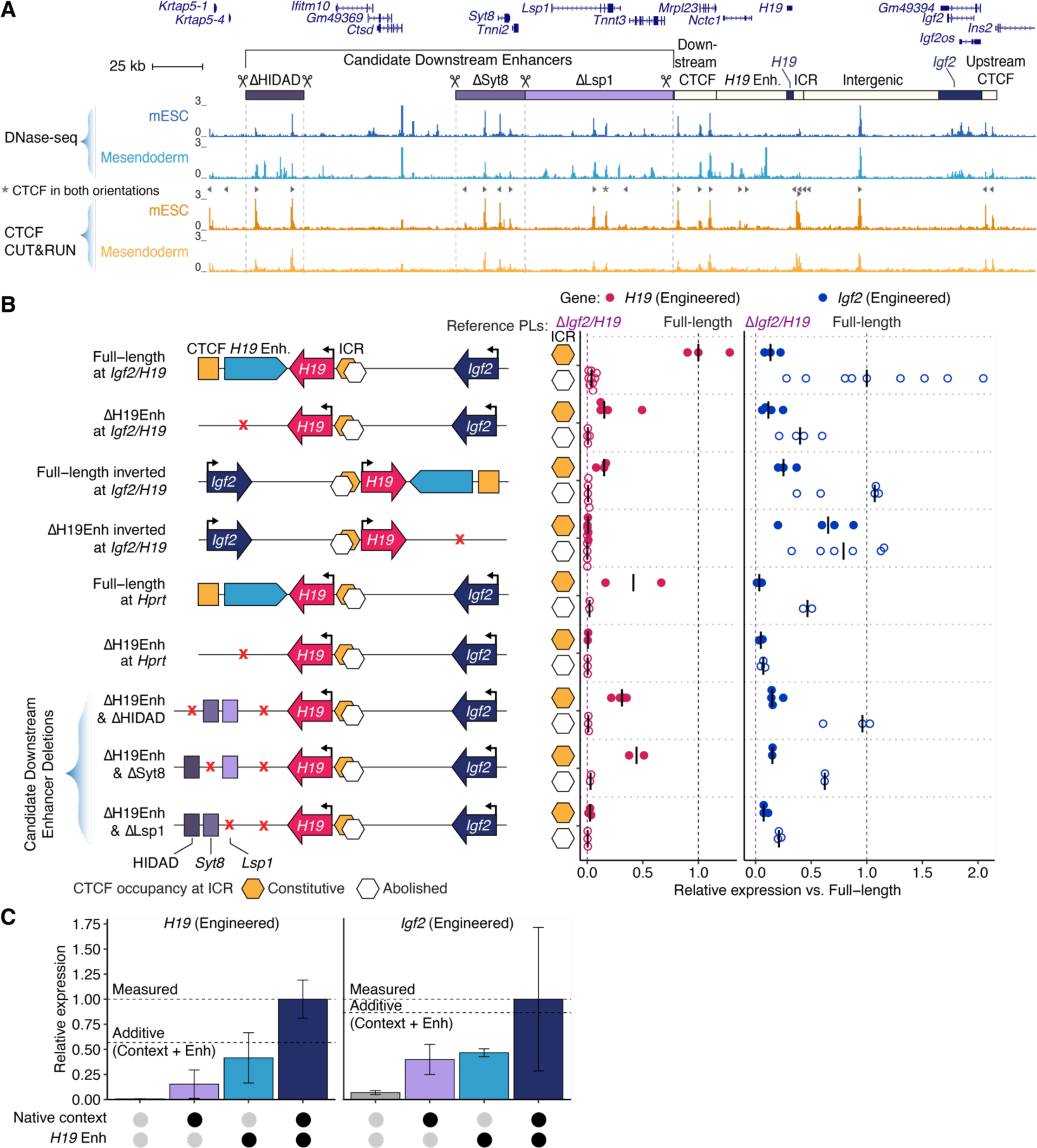
Identification of long-range regulatory elements at the *Igf2/H19* locus. (**A**) The *Igf2/H19* locus and extended downstream region showing DNase-seq and CTCF CUT&RUN in mESC and differentiated mesendodermal cells. Triangles indicate CTCF recognition sequence orientation. “Full-length” and “ΔH19Enh at *Igf2/H19*” are repeated from Figure 1E for reference. (**B**) Expression analysis in mesendodermal cells of *Igf2/H19* payloads when delivered to the *Igf2/H19* locus, to a genomic safe harbor locus (*Hprt*), or to *Igf2/H19* locus in conjunction with distal candidate enhancers deletions. Each point represents the expression of the engineered *H19* or *Igf2* in an independent clone using allele-specific qRT-PCR with bars indicating the median. First two rows are from Figure 1E. Expression was normalized to *Gapdh* and then scaled between 0 (Δ*Igf2/H19* locus) and 1 (“Full-length” rescue payloads), normalizing *H19* to the “Full-length” payload with constitutive CTCF occupancy, and *Igf2* to the “Full-length” payload with abolished CTCF occupancy, which are also represented as dashed lines. Payloads with both constitutive CTCF occupancy (orange hexagon) or abolished CTCF occupancy (empty hexagon) at the ICR were assessed. (**C**) Contribution of the *H19* enhancers and the native context in the expression levels of *H19* and *Igf2* genes in the payloads with constitutive CTCF occupancy or abolished CTCF occupancy, respectively. The presence (black dots) and absence (gray dots) of each element is indicated. Bars indicate median and error bars represent the interquartile range. Dashed horizontal lines indicate the expected transcriptional output of adding both the enhancers and the surrounding context (bottom), versus the measured transcriptional output (top). Source data are from (**B**).

As a complementary test of the reliance of *Igf2/H19* expression on their broader genomic environment, we relocated *Igf2/H19* payloads to a completely different genomic location. The *Hprt* locus is often used as a safe harbor for ectopic transgene expression in a permissive genomic context across many cell types. We thus started from a LP-*Hprt* mESC line, wherein a Big-IN landing pad replaces the *Hprt* locus in chromosome X in a male mESC hybrid cell line (Camellato et al. 2024). We additionally engineered a CRISPR/Cas9-mediated deletion of the B6 allele of the full *Igf2/H19* locus in these mESC (LP-*Hprt*-Δ*Igf2/H19* cells) to maintain a single endogenous copy of *Igf2* and *H19* in addition to the delivered payload (**Figure S11A**). When the “Full-length” payloads were delivered ectopically to the *Hprt* locus, not only were *Igf2* and *H19* expressed, but also the ICR-regulated gene expression pattern was correctly recapitulated. However, both *Igf2* and *H19* expression was reduced by 50% as compared to the endogenous locus. Strikingly, *Igf2* and *H19* expression from the *Hprt* locus was completely ablated in “ΔH19Enh at *Hprt*” payloads lacking the *H19* enhancers (**Figure 2B**). This suggests that *Igf2/H19* expression can be deconstructed into two parts: (i) a well-known ICR-regulated contribution of the *H19* enhancers; and (ii) a newly identified independent contribution of the surrounding genomic environment.

Based on these results, we examined three enhancer clusters downstream of *Igf2* and *H19* showing DNase-seq activity in mesendodermal cells (**Figure 2A**) (Llères et al. 2019). All three clusters are contained within the same topological associated domain (TAD) in mESCs as the *Igf2* and *H19* genes (Llères et al. 2019). Three of these regions have been shown to form an allelic sub-TAD structure with *Igf2/H19* genes by 4C-seq and DNA FISH, and were named HIDAD (*H19*/*Igf2* Distal Anchor Domain), *Syt8*, and *Lsp1* (**Figure 2A**) (Llères et al. 2019). We engineered three derivative landing pad mESC lines, each with a CRISPR/Cas9-mediated deletion of one of these regions. We then delivered *Igf2/H19* payloads lacking the *H19* enhancer cluster to each of these landing pad cell lines. “ΔH19Enh” payloads delivered to ΔHIDAD and ΔSyt8 landing pad mESC lines (“ΔHIDAD & ΔH19Enh” and “ΔSyt8 & ΔH19Enh” respectively) showed no difference in expression relative to their delivery to LP-*Igf2/H19* mESCs, while delivery to ΔLsp1 landing pad mESCs (“ΔLsp1 & ΔH19Enh”) showed negligible *H19* expression and reduced *Igf2* expression (**Figure 2B**). This implicates the *Lsp1* enhancer cluster in the regulation of *H19*, and to a lower extent of *Igf2.* We observed a synergistic cooperation between the *H19* enhancers and the surrounding genomic context, with the expression of both genes being greater than the sum of the individual activities (**Figure 2C**). These results highlight the contribution of the surrounding genomic environment to the *Igf2/H19* locus regulation, and demonstrate the power of delivery at defined ectopic sites to establish sufficiency.

### Enhancer sufficiency and robustness can be disrupted in different genomic contexts

We then asked whether the *H19* enhancer cluster demonstrated context sensitivity similar to that of the entire *Igf2/H19* locus. The *H19* enhancer cluster is a composite element composed of 11 DHSs covering 55 kb. We compared its function with the *Sox2* locus control region (LCR) which, similar to the *H19* enhancers, consists of 10 DHSs across 41 kb located also approximately 100 kb away from its target gene (Zhou et al. 2014; Li et al. 2014; Brosh et al. 2023) (**Figure 3A**). In contrast to the *H19* enhancers, the *Sox2* LCR displays strong activity in mESCs, which is retained into early mesendoderm differentiation (**Figure S3**). We therefore exchanged enhancer clusters among these two loci to establish the extent to which their function was tied to other elements of their surrounding locus.

**Figure 3.**
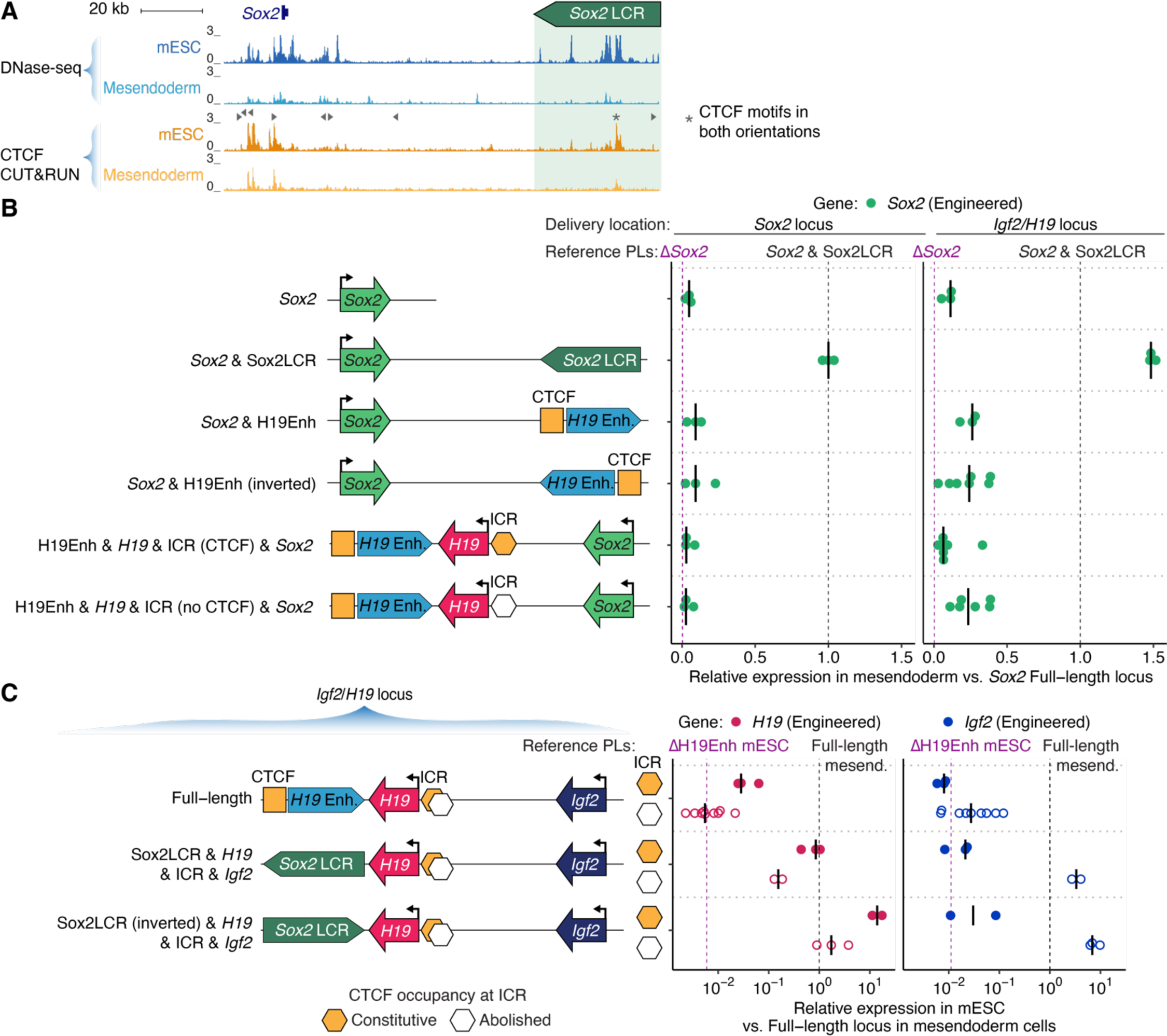
Enhancer interchangeability at the *Igf2*/*H19* and *Sox2* loci. (**A**) *Sox2* and locus control region (LCR) showing DNase-seq and CTCF CUT&RUN in mESC and mesendodermal cells. Triangles indicate CTCF recognition sequence orientation. (**B-C**) Points represent independently engineered clones and bars indicate the median (**B**) Expression analysis in differentiated mesendodermal cells harboring payloads including the wild-type *Sox2* locus, the *Sox2* locus with the LCR replaced by the *H19* enhancer cluster, and the *Igf2*/*H19* locus with the *Igf2* gene and promoter replaced by the *Sox2* gene and promoter. Each payload was delivered both to the *Sox2* locus (left) and *Igf2/H19* locus (right). Expression of the engineered *Sox2* allele was normalized to the wild-type CAST allele and then scaled between 0 (Δ*Sox2* locus) to 1 (“*Sox2* & Sox2 LCR”), which are also represented as dashed lines. (**C**) *Igf2* and *H19* expression analysis in mESC engineered with a “Full-length” *Igf2/H19* rescue payload, or payloads wherein the *H19* enhancer was replaced by the *Sox2* LCR in either orientation. Expression was normalized to *Gapdh* and then scaled between 0 (Δ*Igf2/H19* locus) and 1 (“Full-length”), normalizing *H19* to the “Full-length” payload with constitutive CTCF occupancy, and *Igf2* to the “Full-length” payload with abolished CTCF occupancy. Payloads with both constitutive CTCF occupancy (orange hexagon) or abolished CTCF occupancy (empty hexagon) at the ICR were assessed. The dashed lines indicate basal expression of “ΔH19Enh” in mESCs and expression of “Full-length” *Igf2/H19* rescue payload in differentiated mesendodermal cells.

We employed LP-*Sox2* and LP-LCR mESCs harboring Big-IN landing pads replacing the full 143-kb *Sox2* locus or just the *Sox2* LCR, respectively (Brosh et al. 2023). We quantified the expression of the engineered *Sox2* allele on a scale of 0 (ΔSox2) to 1 (“*Sox2* & Sox2LCR”) using qRT-PCR (**Figure 3B**). We assembled payloads comprising just the *H19* enhancers in both orientations (**Figure 3B**). When the *H19* enhancers were delivered to LP-LCR mESCs replacing the *Sox2* LCR, *Sox2* expression remained very low even upon differentiation to mesendodermal cells (**Figure 3B**, left panel, “*Sox2* & H19Enh” and “*Sox2* & H19Enh (inv)”). To test whether the *H19* enhancers relied upon other regulatory elements within the 157-kb payload boundaries, we edited the “Full-length” *Igf2/H19* payload in yeast to replace the *Igf2* gene and its promoter with the *Sox2* gene and promoter (**Figure S12**), and delivered it to LP-*Sox2* mESCs replacing the whole *Sox2* locus (**Figure 3B**, “H19Enh & *H19* & ICR (CTCF) & *Sox2*” and “H19Enh & *H19* & ICR (no CTCF) & *Sox2*”). These too showed very low *Sox2* expression (**Figure 3B**, left panel), suggesting that the *H19* enhancers were either incompatible with the *Sox2* promoter or lacked key elements from outside the *Igf2*/*H19* “Full-length” payloads.

To investigate whether the *H19* enhancers relied on genomic elements outside the boundaries of our 157-kb payload, we delivered the same payloads at the *Igf2/H19* locus (**Figure 3B**, right panel). Starting from LP-*Igf2/H19* mESCs, we further engineered a CRISPR/Cas9-mediated deletion of the endogenous B6 *Sox2* allele (LP-*Igf2/H19*-Δ*Sox2* cells, **Figure S11B**) to ensure only two copies of *Sox2* were present after payload delivery. A payload with the full *Sox2* locus including its native LCR (“*Sox2* & Sox2LCR”) resulted in a 1.5-fold increase in *Sox2* expression when delivered to the *Igf2/H19* locus, relative to its endogenous location. In comparison, both payloads wherein the *H19* enhancers replaced the *Sox2* LCR (“*Sox2* & H19Enh” and “*Sox2* & H19Enh (inv)”, right panel) showed a 2.5-fold increase in *Sox2* expression when delivered to the *Igf2/H19* locus compared to at the *Sox2* locus. Notably, the intervening sequence between the *H19* enhancers and *Igf2* was not required to activate *Sox2*, as the *H19* enhancers drove similar *Sox2* expression in payloads derived from the *Sox2* locus (“*Sox2* & H19Enh” and “*Sox2* & H19Enh (inv)”) and from the *Igf2/H19* locus (H19Enh & *H19* & ICR (CTCF/no CTCF) & *Sox2*”). Finally, the *H19* enhancers remained sensitive to CTCF occupancy at the intervening ICR, activating the *Sox2* promoter only in absence of CTCF (**Figure 3B**).

We next performed the cognate swap, replacing the *H19* enhancers with the 41-kb *Sox2* LCR in both orientations, yielding “Sox2LCR & *H19* & ICR & *Igf2*” and “Sox2LCR (inv) & *H19* & ICR & *Igf2*” payloads. We profiled *Igf2* and *H19* expression in mESCs and scaled expression to that of the “Full-length” payloads in mesendodermal cells for comparison. This showed that the *Sox2* LCR robustly boosted the expression of both *Igf2* and *H19* in undifferentiated cells relative to their native state (**Figure 3C, Figure S3**). The convergent orientation of the *Sox2* LCR (i.e. with CTCF sites oriented towards the target genes) yielded increased expression of both *Igf2* and *H19* genes relative to the *Sox2* LCR in the divergent orientation (**Figure 3C**), consistent with observations at the endogenous *Sox2* locus (de Wit et al. 2015; Brosh et al. 2023). In either orientation, the *Sox2* LCR activated *Igf2* almost exclusively in the absence of CTCF binding at the ICR, matching the behavior of the endogenous *H19* enhancer cluster (**Figure 3C**). But, in contrast to the *H19* enhancers, the *Sox2* LCR upregulated *H19* expression in the absence of CTCF occupancy. Put another way, CTCF occupancy at the ICR resulted in only a 5-8-fold difference in *H19* expression, compared to a >100-fold difference in *Igf2* expression (**Table 1**). When linked to the *Sox2* LCR instead of the *H19* enhancers, the relative effect of the ICR was weaker on *H19* expression but stronger on *Igf2* expression (**Table 1**). Thus, the *Sox2* LCR did not fully recapitulate the activity of the endogenous regulation of the *H19* enhancer cluster. Understanding this promiscuity in more detail may provide important clues to the function of the endogenous ICR and the sequence determinants of enhancer selectivity.

**Table 1.**
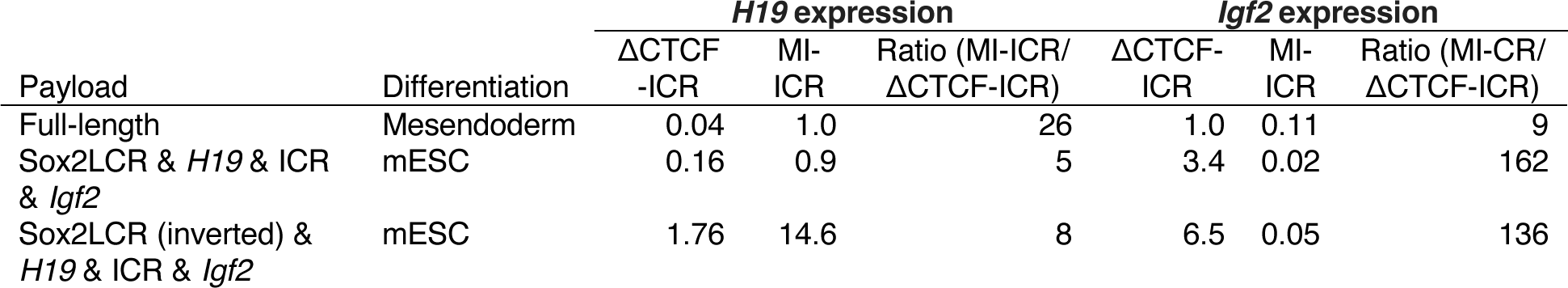
Comparison of ICR function when linked to *H19* enhancers and the *Sox2* LCR. Comparison of *H19* and *Igf2* expression for pairs of payloads with either constitutive CTCF occupancy (MI-ICR) or abolished CTCF occupancy (ΔCTCF-ICR). Expression was normalized to *Gapdh* and then scaled between 0 (Δ*Igf2/H19* locus) and 1 (“Full-length”), normalizing *H19* to the “Full-length” payload with constitutive CTCF occupancy, and *Igf2* to the “Full-length” payload with abolished CTCF occupancy.

### Characteristic patterns of coordinate DHS actuation across cell and tissue types

Regulatory elements operate within complex neighborhoods, including extended regions of coordinated transcriptional programs as well as examples of genes with distinct expression patterns relative to their neighbors (Pachano et al. 2022). The contrasting function of the *Sox2* LCR and *H19* enhancer cluster in ectopic contexts suggests they may have different relationships with their native genomic context. We analyzed cell-type selective accessibility at the *Hprt*, *Sox2*, and *Igf2*/*H19* loci using DNase-seq data from 45 representative cell and tissue types from the mouse ENCODE project (Vierstra et al. 2014). *Hprt* showed accessibility in all cell and tissue types, befitting its status as a housekeeping gene (**Figure S13A**). In contrast, the *Sox2* LCR was more specific, while the *Sox2*-proximal DHSs and those at *Igf2*/*H19* showed broader tissue-selective accessibility profiles. We further observed stereotypical differences in the pattern of coordinated DHS activity across tissue types surrounding each of the delivery sites (**Figure 4A)**. The *Sox2* LCR was active specifically in mESC, while the rest of the locus harbored a series of more broadly active enhancers. This is consistent with the observation that the LCR is required for *Sox2* expression in mESC but becomes silent and dispensable upon differentiation to neuroectoderm (Zhou et al. 2014). A broader pattern of coordinated activity extended across the *Igf2* and *H19* genes and partially into *Lsp1*. In contrast, the *Hprt* locus was sparsely populated by mostly constitutive DHSs, most prominently the DHSs at the *Hprt* and *Phf6* promoters. The loci also differed in terms of gene and DHS density: *Sox2* lies within a gene desert, *Igf2* and *H19* lie in a region rich with other genes and DHSs, and *Hprt* is surrounded by an intermediate number of genes (**Table 2**).

**Figure 4.**
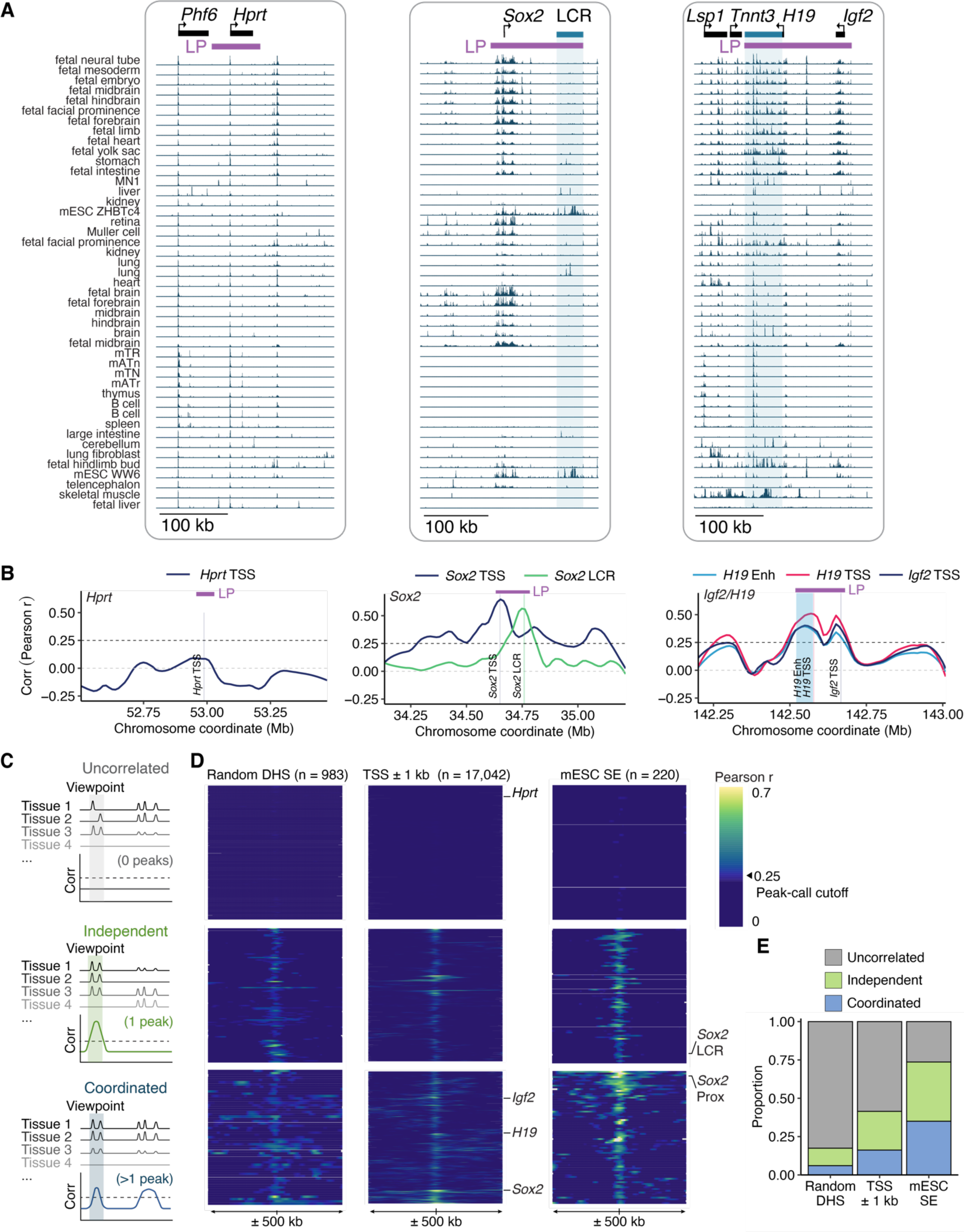
Cross-cell type accessibility patterns reflect locus function. (**A**) Accessibility patterns at the *Hprt*, *Sox2* and *Igf2/H19* loci across 45 cell and tissue types, ordered by hierarchical clustering of genomic DHSs as in (Halow et al. 2021). Shading represents the *Sox2* LCR and *H19* enhancers. (**B**) Correlation of DHS accessibility across *Hprt*, *Sox2* and *Igf2/H19* loci. Each line represents the Pearson correlation coefficient (r) between the DHS located at the viewpoint with every other DHS in a 1-Mb window. Viewpoints are shaded in each plot. (**C**) Schematic representation of uncorrelated, independent and coordinated DHS clusters. Multiple regions of high Pearson r values indicate coordinated DHS activity potentially due to synchronized activation in specific tissues. Corr: Pearson correlation coefficient. (**D**) Heatmaps showing the Pearson correlation coefficient (r) between the DHS located at the viewpoint and the surrounding DHS within a 1-Mb window. Three different sets of viewpoints are represented: random DHSs, promoters of protein coding genes (±1 kb from TSS), and mESC super-enhancers (SE) (Whyte et al. 2013); each row portrays the region surrounding a different viewpoint. Rows are not shown to scale. DHS correlation patterns were grouped into uncorrelated, independent and coordinated based on the number of peaks exceeding *r* > 0.25. (**E**) Representation of DHSs correlation patterns for various genomic features.

**Table 2.**
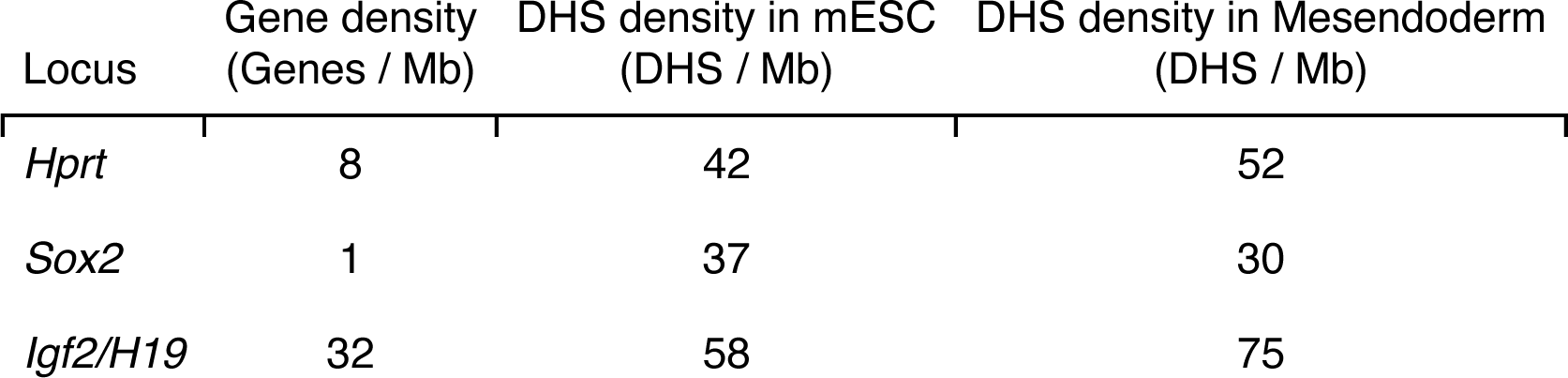
Regulatory environment complexity. Gene and DHS density was calculated by counting the number of genes or DHS hotspots (FDR < 0.05), respectively, in a window of 1 Mb sequence centered on each locus. Genes include all validated (level 1) or manually annotated (level 2) Gencode VM23 protein coding transcripts.

| Locus | Gene density (Genes / Mb) | DHS density in mESC (DHS / Mb) | DHS density in Mesendoderm (DHS / Mb) |
| --- | --- | --- | --- |
| *Hprt* | 8 | 42 | 52 |
| *Sox2* | 1 | 37 | 30 |
| *Igf2/H19* | 32 | 58 | 75 |

Regulatory interaction can be reflected in correlated actuation of regulatory DNA across cell types (Thurman et al. 2012). We thus computed correlations in accessibility across 45 cell and tissue types between the *Hprt*, *Sox2,* and *Igf2*/*H19* loci and their surrounding 1 Mb (**Figure 4B**, **Figure S13B-D**). While the *Hprt* transcription start site (TSS) showed no correlation with surrounding DHSs, the *Sox2* LCR DHSs showed correlation only amongst themselves. Finally, the *Sox2* TSS as well as the *Igf2* TSS, *H19* TSS, and *H19* enhancers showed tight correlation with additional surrounding regions. We defined these three patterns as uncorrelated, independent, and coordinated activity, respectively, and analyzed their representation more broadly across the genome (**Figure 4C-D**). A random sample of genomic DHSs exhibited uncorrelated activity (**Figure 4E**). Focusing on gene promoters (±1 kb from TSS) showed stronger correlation with surrounding regions. Finally, mESC super-enhancers (Whyte et al. 2013) exhibited the highest rate of independent and coordinated activity, with only 27% being uncorrelated. These cross-cell type actuation patterns may provide a key feature of genomic context at tissue-selective genes and may distinguish “independent” from “context-dependent” enhancer clusters.

## DISCUSSION

Our work shows that composite regulatory elements such as enhancer clusters recapitulate their characteristic function ectopically with degraded fidelity relative to their full native context. The *Sox2* LCR at first glance manifests the autonomy and independence implied by the classical definition of an enhancer (Banerji et al. 1981), in that it drives robust expression at the *Igf2*/*H19* locus. Yet its influence on *H19* expression differs from the endogenous *H19* enhancers in that the *Sox2* LCR partially overrides the ICR, upregulating *H19* even in the absence of CTCF occupancy. The *H19* enhancers and ICR both harbor multiple CTCF sites convergent on *H19*, and the *Sox2* LCR harbors CTCF sites pointing towards *Sox2*. Inverting the ICR has been shown to lower the range of expression between the paternal and maternal alleles (Matsuzaki et al. 2021), and we show that inversion of the *Sox2* LCR in the context of the ICR yields reduced expression, especially for *H19* (**Figure 3C**). This suggests a CTCF-mediated link between the ICR and the enhancer cluster it modulates, although we note that the link between LCR orientation and expression at *Sox2* is not due solely to CTCF (de Wit et al. 2015; Brosh et al. 2023). It is possible that CTCF sites underpin the robustness of the *Sox2* LCR across genomic positions or its interaction with the ICR (Ribeiro-Dos-Santos et al. 2022), but this does not explain why the CTCF sites in the *H19* enhancers behave differently. Thus more work is needed to further dissect whether there are specific sequence determinants within the *Sox2* LCR (i.e. DHSs or TF recognition sequences) responsible for such distinctive activity, or if its autonomy simply derives from its strength.

The *H19* enhancers themselves are highly subordinated to their enclosing genomic context, and show reduced activity upon any perturbation in the surrounding architecture, such as *Lsp1* deletion, delivery to an ectopic locus, or linkage to a heterologous promoter. Our work thus shows complex regulatory elements are subject to context sensitivity, in contrast to the view that such elements can exert dominant effects. Future research will be needed to establish the range at which this context requirement operates, and whether this requirement is tied to a single element, or reflects the contributions of many. For the *H19* enhancers, our data show that the necessary element(s) lie outside the 157-kb *Igf2*/*H19* locus. It is unclear whether these partners resemble the context-dependent or ‘facilitator’ DHSs previously reported (Blayney et al. 2023; Brosh et al. 2023). Or, there might be repressive elements outside the 143-kb *Sox2* locus that suppress the *H19* enhancers. We also note that while *Sox2* and *Igf2* are at comparable distances to their native enhancer clusters, *H19* lies much closer, and it is possible that the *H19* enhancers would be able to activate a target gene ectopically at closer range. Finally, robust domain formation relies on CTCF sites throughout a locus (Despang et al. 2019). It is possible that the *Lsp1* enhancers themselves underpin the synergy with the *H19* enhancers. Additionally, while the deletion of the *Syt8* CTCF cluster did not affect *H19* expression, it slightly increased *Igf2* expression. Similarly, HIDAD includes a pair of CTCF sites that have been shown to interact with elements within the *Igf2*/*H19* locus (Rao et al. 2014; Ye and Ma 2020; Richer et al. 2023), and its deletion increased *Igf2* expression. Thus it is possible the CTCF sites within these regions underpin the overall genomic organization of the locus and their deletion could sensitize it to further disruption.

Our analysis of cross-cell type correlation in regulatory DNA actuation suggests a broad influence of genomic context in potentiating enhancer cluster function. While there is little evidence for strict genomic regulons in complex organisms, domains of coordinated expression have been observed in yeast (Cohen et al. 2000) and fly (Rubin and Green 2013). Mammalian genomes demonstrate a certain compartmentalization of function, in that housekeeping and disease-associated genes are unevenly distributed genome-wide (Lercher et al. 2002; Muro et al. 2019), and TADs correlate with tissue-selective expression patterns (Symmons et al. 2014). But the strength, generalizability, and sequence determinants of these architectural context effects have been difficult to clarify across mammalian genomes. Our analysis partitions genomic enhancer clusters into two classes – Independent and Coordinated – which differ in their interaction with genomic context. This suggests that the coordinated tissue-specific accessibility and expression patterns observed at key developmental loci (Bolt and Duboule 2020) might be required to reinforce the intrinsic propensities of regulatory elements therein. This is consistent with the β-globin genes, whose promoters add developmental specificity to the activation provided by the LCR (Bender et al. 2000). The large gene deserts surrounding tissue-selective genes (e.g. *Sox2* and *Hbb*) may be required to insulate unrelated surrounding genes from this context effect, and might derive mechanistically from a reliance on cohesin for longer-range contacts for tissue-specific genes (Kane et al. 2022; Rinzema et al. 2022) or models invoking diffusion of regulatory factors (Karr et al. 2022). We note that while enhancers showing correlated cross-cell type accessibility patterns have appeared as redundant in deletion analyses (Hong et al. 2008; Frankel et al. 2010; Osterwalder et al. 2018; Hörnblad et al. 2021), our sufficiency analysis suggests that these broader activity patterns can be required for proper function.

In this study, we showcase the potential of synthetic regulatory genomics to characterize and interrelate multiple regulatory elements, to interpret complex loci, and to gain new insights into transcriptional regulation. This approach can be easily applied to characterize many other genomic regions whose size, complexity, and/or allele-specific regulation challenge the limited scope of genome editing tools currently in wide use. We expect that dissecting specific sequence elements responsible for novel activities of synthetic combinations of regulatory elements will lead to generalizable models of locus architecture and gene regulation.

### Limitations of the study

Our study is based on measurement of an expression phenotype after differentiation. While we have performed measurements across replicate clones, our results might still be affected by the reproducibility of differentiation, or differences between clones in general, including variability between payloads delivered to different landing pad parental lines. While we investigated activity in mESCs and differentiated mesendodermal cells, accessibility at the *H19*/*Igf2* locus shows a variety of activity patterns across different tissues (**Figure 4A**), and their regulation might differ from the contexts studied here. We have developed a genetic strategy for manipulating CTCF occupancy at the ICR in cell culture, but it is possible that locus function behaves differently upon passage through the germline. Finally, we only investigated two enhancer clusters and others may behave differently.

## ACKNOWLEDGEMENTS

We thank Brendan Camellato for the gift of pDONOR∼Neo∼EGFP, and Raven D. Luther for technical assistance with sequencing. This work was partially funded by National Institutes of Health (NIH) grants RM1HG009491 (to J.D.B.) and R35GM119703 (to M.T.M.).

## AUTHOR CONTRIBUTIONS

R.O., M.S.H., and M.T.M. designed experiments. R.O., G.E., and R.B. performed experiments. W.Z. and Y.Z. assembled DNA payloads. G.E., H.J.A., and E.H. performed capture and sequencing. R.O. and A.M.R.-d.-S. performed computational analyses. R.O. and M.T.M. wrote the manuscript. All authors edited the manuscript. M.T.M. and J.D.B. supervised work.

## STAR METHODS

### KEY RESOURCES TABLE

| REAGENT or RESOURCE | SOURCE | IDENTIFIER |
| --- | --- | --- |
| **Bacterial and virus strains** |  |  |
| Parental *S. cerevisiae* strain | Brachmann et al., 1998(Brachmann et al. 1998) | BY4741 |
| Payload *S. cerevisiae* strain | This paper | Table S1 |
| *E. Coli* TransforMax EPI300 | Lucigen | EC300150 |
| **Chemicals, peptides, and recombinant proteins** |  |  |
| 0.1% gelatin | EMD Millipore | ES006-B |
| 2-mercaptoethanol | Sigma | M3148 |
| Advanced DMEM/F-12 | ThermoFisher | 12634010 |
| BbsI-HF | NEB | R3539L |
| Blasticidin | ThermoFisher | A1113902 |
| BsaI-HF V2 | NEB | R3733S |
| CHIR99021 | R&D Systems | 4423 |
| CopyControl Induction Solution | Lucigen | CCIS125 |
| DNA endonuclease DNase I | Sigma | D4527-20KU |
| Esp3I | NEB | R0734S |
| FBS | BenchMark | 100-106 |
| ganciclovir | Sigma | PHR1593 |
| geneticin G418 | ThermoFisher | 10131035 |
| GlutaMAX | ThermoFisher | 35050061 |
| GoTaq Green | Promega | M7123 |
| KAPA 2x Hi-Fi Hotstart Readymix | Roche | 07958935001 |
| KnockOut DMEM | ThermoFisher | 10829018 |
| LIF | Sigma | ESG1107 |
| MEM nonessential amino acids | ThermoFisher | 11140050 |
| N2 Supplement | ThermoFisher | 17502048 |
| Neurobasal | ThermoFisher | 21103049 |
| nucleosides | EMD Millipore | ES008-D |
| PD0325901 | Sigma | PZ0162 |
| Pen-Strep | ThermoFisher | 15140122 |
| proaerolysin | Aerohead Scientific | n/a |
| Proteinase K | ThermoFisher | EO0491 |
| puromycin | ThermoFisher | A1113803 |
| RNAse A | Sigma | R4642-50MG |
| Sodium Pyruvate | ThermoFisher | 11360070 |
| TrypLE-Select | ThermoFisher | 12563011 |
| Y-27632 ROCK inhibitor | Cell Signaling Technology | 13624S |
| **Antibodies** |  |  |
| CTCF antibody | Cell Signaling | 2899S |
| **Critical commercial assays** |  |  |
| CUT&RUN Assay Kit | Cell Signaling | 86652 |
| DNeasy Blood & Tissue kit | QIAGEN | 69506 |
| KAPA SYBR FAST | Kapa Biosystems | KK4610 |
| Multiscribe High-Capacity cDNA Reverse Transcription Kit | ThermoFisher | 4368814 |
| NEBNext Poly(A) mRNA Magnetic Isolation Module | NEB | E7490 |
| NEBNext Ultra II Directional RNA Library Prep Kit | NEB | E7760S |
| Nucleobond XtraBAC kit | Takara | 740436 |
| Turbo DNA-free kit | ThermoFisher | AM1907 |
| Zymo yeast miniprep I protocol | Zymo Research | D2001 |
| ZymoPURE maxiprep kit | Zymo Research | D4203 |
| Zyppy Plasmid Miniprep Kit | Zymo Research | D4020 |
| **Deposited data** |  |  |
| Sequencing data | This paper | GSE241708 |
| **Experimental models: Cell lines** |  |  |
| Parental C57BL6/6J × CAST/EiJ (BL6xCAST) mESCs | Eckersley-Maslin et al., 2014(Eckersley-Maslin et al. 2014) | Clone 4 |
| C57BL6/6J × CAST/EiJ (BL6xCAST) Landing Pad and Payload mESC clones | This paper | Table S3 |
| **Oligonucleotides** |  |  |
| Cloning primers | This paper | Table S6 |
| Genotyping primers | This paper | Table S6 |
| qRT-PCR primers | This paper | Table S6 |
| Synthetic fragments | This paper | Table S6 |
| **Recombinant DNA** |  |  |
| Payloads | This paper | Table S1, Data S1 |
| pCAG-iCre | Addgene | 89573 |
| pDONOR-tagBFP-PSM-EGFP | Addgene | 100603 |
| pLM1110 | Addgene | 168460 |
| pLM1110_URA3 | This paper | n/a |
| pLP-PIGA2 (pLP300) | Addgene | 168462 |
| pYTK-Cas9 | Zhao et al., 2022(Zhao et al. 2023) | n/a |
| **Software and algorithms** |  |  |
| Sequencing processing pipeline | This paper | TBD |
| BWA | Li and Durbin 2009(Li and Durbin 2009) | https://github.com/lh3/bwa |
| bcftools | Bonfield et al.,<br>2021(Bonfield et al.<br>2021) | https://github.com/samtools/bcftools |
| BEDOPS | Neph et al.,<br>2012(Neph et al.<br>2012) | https://bedops.readthedocs.io/en/latest/ |
| R | R Core Team 2018(R<br>Core Team 2018) | https://www.r-project.org |
| RSEM | Li and Dewery<br>2011(Li and Dewey<br>2011) | https://github.com/deweylab/RSEM |
| STAR | Dobin et al.<br>2013(Dobin et al.<br>2013) | https://github.com/alexdobin/STAR |

### RESOURCE AVAILABILITY

#### Lead contact

Further information and requests for resources and reagents should be directed to and will be fulfilled by the Lead Contact, Matthew T. Maurano.

#### Materials availability

All unique/stable reagents generated in this study are available upon request from the Lead Contact with a completed Materials Transfer Agreement.

#### Data and code availability

Sequencing data have been deposited in the Gene Expression Omnibus (GEO) database, https://www.ncbi.nlm.nih.gov/geo (accession no. GSE241708). DNase-seq data were obtained from https://www.encodeproject.org for ES_E14 (ENCLB890BTE, DS21450). ChIP-nexus (Avsec et al. 2021) for Nanog (GSM4072778) and RNA-seq (GSE222038 samples GSM6912138-GSM6912143) for mESC were obtained from the GEO repository.

### EXPERIMENTAL MODEL AND SUBJECT DETAILS

#### Landing Pads and CRISPR/Cas9 plasmids for genome integration

pLP-PIGA2 (pLP300) (Addgene 168462) (**Figure 1C** and **Figure S1A**) was described previously (Brosh et al. 2021) and harbors a pEF1α-PuroR-P2A-hmPIGA-P2A-mScarlet-EIF1pA cassette flanked by LoxM and LoxP sites, as well as a pPGK1-ΔTK-SV40pA backbone counter-selectable marker cassette. To target landing pads to specific genomic loci, guide RNAs (gRNAs) (**Table S6**) were cloned into pSpCas9(BB)-2A-Puro (pCas9-Puro, Addgene 62988) using BbsI Golden Gate reactions as described (Ran et al. 2013). Landing pad homology arms (HAs) corresponding to the sequence flanking genomic Cas9 cut sites were amplified from BACs (**Table S7**). HAs were cloned to flank the LoxM and LoxP sites in pLP-PIGA2 using a BsaI Golden Gate reaction (**Table S6**). DNA suitable for transfection was prepped using the Zymo-PURE maxiprep kit (Zymo Research D4203) according to the manufacturer’s protocol.

#### Payload assembly

All payloads were assembled into Yeast Assemblon Vector (YAV) pLM1110 (Brosh et al. 2021) (Addgene #168460). This is a multifunctional vector supporting (i) low-copy DNA propagation, selection (*LEU2*), and efficient homology-dependent recombination in yeast; (ii) low-copy propagation, selection (KanR), and copy number induction in TransforMax EPI300 *E. coli* (Lucigen EC300150) (Wild et al. 2002); and (iii) transient visualization and selection (eGFP-BSD, enhanced green fluorescent protein - Blasticidin-S deaminase) in mammalian cells (Brosh et al. 2021).

Three different strategies were used to assemble payloads into pLM1110 (**Table S1**), using different primers, gRNAs, and synthetic DNA fragments (**Table S6**):

##### (1) BAC fragment assembly in yeast

Desired segments of *Igf2/H19* BAC RP23-50N22 (**Table S7**) were released by an *in vitro* CRISPR/Cas9 digestion using a pair of synthetic gRNAs and recombinant Cas9 and assembled into BsaI-digested pLM1110 as previously described (Brosh et al. 2021).

##### (2) CRISPR/Cas9 Engineering of EPisomes in Yeast (CREEPY)

Existing payloads were modified in yeast, as in Zhao et al. (Zhao et al. 2023), using CRISPR/Cas9 and synthetic linker fragment(s) (IDT) to mediate sequence deletions, insertions, or inversions. Yeast cells carrying a parental payload were transformed with pYTK-Cas9 plasmids (single-plasmid system co-expressing a yeast-codon optimized Cas9, the targeting gRNA and a *HIS3* marker) together with 100 ng of the synthetic linker fragment(s) (**Table S1**), and selected for a Leu^+^/His^+^ phenotype. gRNAs (**Table S6**) were cloned into pYTK-Cas9 using a Esp3I Golden Gate reaction.

##### (3) Payload inversion

To deliver payloads in an inverted orientation, the flanking Lox sites were swapped in yeast. Source constructs were first digested using I-SceI, releasing the payload sequence from the backbone including the Lox sites. This digestion product was then co-transformed into yeast cells with I-SceI-digested pLM1110_URA3 (a derivative of pLM1110 carrying a *URA3* marker instead of *LEU2*) and terminal linker fragments including the repositioned Lox sites (400-500 bp gBlocks, IDT) (**Table S1**, **Table S6**) to enable homologous recombination-dependent assembly. Yeast were then selected for a Ura^+^ phenotype, depleting the wild-type Leu^+^ recombinants.

#### Yeast transformation, payload validation, and transfer to *E. coli*

The following steps were performed as previously described (Brosh et al. 2023). Specifically, yeast transformations were performed using the Lithium acetate method (Gietz and Schiestl 2007). For screening of correct clones, DNA was isolated from individual yeast colonies by resuspension in 10-40 µL of 20 mM NaOH and boiling for 3 cycles of 95 °C for 3 min and 4°C for 1 min. 2 µL of yeast lysate was used as a template in a 10 µL GoTaq Green reaction (Promega M7123) with 0.25 µM of primers. Clones were screened for the intended payload structure using primers that target newly formed junctions (positive screen), and in some cases using primers that target regions that are present in the parental plasmid but absent in the intended plasmid (negative screen). Assembled payloads were isolated from candidate yeast clones using the Zymo yeast miniprep I protocol (Zymo Research D2001) and transformed into TransforMax EPI300 *E. coli* cells by electroporation using the manufacturer’s protocol.

#### Payload DNA screening and preparation

Payload DNA was prepped and verified as previously described (Brosh et al. 2023). Briefly, for initial verification, TransforMax EPI300 *E. coli* colonies were picked into 3-5 mL LB-Kan supplemented with CopyControl Induction Solution (Lucigen CCIS125) and cultured overnight at 30 °C with shaking at 220 RPM. Crude DNA was extracted and payload assembly verification was performed by diagnostic digestion with restriction enzymes, PCR verification of new junctions, and Sanger sequencing, depending on the payload. All assembled payloads were subsequently sequenced to high coverage depth (**Table S8**).

Sequence-verified payload clones were grown overnight in 2.5 mL cultures of LB-Kan at 30 °C with shaking, diluted 1:100 in LB-Kan supplemented with CopyControl Induction Solution, and incubated for an additional 8-16 hours at 30 °C with shaking. For transfection, DNA was purified from induced *E. coli* using the Nucleobond XtraBAC kit (Takara 740436) and preps were stored at 4 °C.

#### mESC culture

mESC were cultured as previously described (Brosh et al. 2023). Specifically, C57BL/6J × CAST/EiJ (B6xCAST) mESCs were cultured on plates coated with 0.1% gelatin (EMD Millipore ES006-B) in 80/20 medium comprising 80% 2i medium and 20% mESC medium. 2i medium contained a 1:1 mixture of Advanced DMEM/F12 (ThermoFisher 12634010) and Neurobasal-A (ThermoFisher 10888022) supplemented with 1% N2 Supplement (ThermoFisher 17502048), 2% B27 Supplement (ThermoFisher 17504044), 1% GlutaMAX (ThermoFisher 35050061), 1% Pen-Strep (ThermoFisher 15140122), 0.1 mM 2-mercaptoethanol (Sigma M3148), 1,250 U/mL LIF (ESGRO ESG1107l), 3 μM CHIR99021 (R&D Systems 4423), and 1 μM PD0325901 (Sigma PZ0162). mESC medium contained knockout DMEM (ThermoFisher 10829018) supplemented with 15% FBS (BenchMark 100-106), 0.1 mM 2-mercaptoethanol, 1% GlutaMAX, 1% MEM nonessential amino acids (ThermoFisher 11140050), 1% nucleosides (EMD Millipore ES008-D), 1% Pen-Strep, and 1,250 U/mL LIF. Cells were grown at 37 °C in a humidified atmosphere of 5% CO2 and passaged twice per week on average.

#### mESC genome engineering

Landing pad integrations were performed using the Neon Transfection System as previously described into B6xCAST ΔPiga mESCs, in which the endogenous Piga gene was deleted (Brosh et al. 2021). LP-PIGA2 integration at the *Igf2/H19* locus (replacing a 157-kb genomic region) was performed using 5 µg of pLP-PIGA2 and 2.5 µg of each pCas9-GFP plasmid expressing B6-specific gRNAs mIgf2_1b and mIgf2_4b, followed by 1 µg/mL puromycin (ThermoFisher A1113803) selection for mESCs harboring a landing pad and 1 µM ganciclovir (GCV, Sigma PHR1593) selection against LP-PIGA2 backbone’s HSV1-ΔTK gene.

The genomic deletion of the *Igf2/H19* B6 allele was performed by replacing the 157-kb genomic region with a selectable module (**Figure S11A**). The donor vector pDONOR∼Neo∼EGFP contained a positive selection cassette expressing eGFP, a Neomycin resistance gene and a negative selection marker expressing tagBFP and was derived from pDONOR-tagBFP-PSM-EGFP (Addgene 100603). Flanking the positive selection cassette, 500 bp homology arms corresponding to the genomic sequence flanking the Cas9 cut sites were cloned. Landing pad cells were further transformed using 5 µg of pDONOR∼Neo∼EGFP vector and 2.5 µg of each pCas9-GFP plasmid expressing B6-specific gRNAs mIgf2_1b and mIgf2_4b, followed by 400 µg/mL Geneticin G418 (Thermo Fisher, 10131035) selection for 4 days.

Genomic deletions of the *Sox2* locus (**Figure S11B**) and the downstream enhancer clusters (*Lsp1*, *Syt8*, and *HIDAD*) were performed by co-transfecting cells with 2.5 μg of each pCas9-GFP and 2.5ul of 100 µM ssODN (IDT) bridging oligo with an overlap of 50 bp on each side of the deletion and modified with 5’ and 3’ Phosphorothioate bonds to prevent degradation. The following gRNAs were used for each deletion: mSox2_5p-1 and mSox2_3p-5 for the *Sox2* locus; mIgf2_26b and mIgf2_27b for the HIDAD DHS cluster; mIgf2_28b and mIgf2_29b for the *Syt8* DHS cluster; mIgf2_29b and mIgf2_30b for the *Lsp1* DHS cluster.

Payload deliveries were performed as previously described (Brosh et al. 2021) with 10 × 10^6^ mESCs, 10 μg payload DNA and 10 μg pCAG-iCre plasmid (Addgene #89573) per transfection. Transfected mESCs were selected with 10 μg/mL blasticidin for 2 days starting day 1 post-transfection and with 2 nM proaerolysin for 2 days starting day 7 post-transfection.

For landing pad integration, payload delivery, and genomic deletions, individual mESC clones were manually picked approximately 10 days post-transfection into gelatinized 96-well plates prefilled with 30 µL TrypLE-Select, which was neutralized with 200 µL mESC medium. Immediately after, 25 µL were transferred into a new plate with 75 µL of 2i media, obtaining two replica plates with 90% and 10% relative densities. Three days later, crude gDNA was extracted from the 90% plate as described (Brosh et al. 2021) and used in PCR genotyping to identify candidate clones, which were then expanded from the 10% density plate for further verification and phenotypic characterization. Genomic DNA was extracted from expanded clones using the DNeasy Blood & Tissue kit (QIAGEN 69506).

### mESC genotyping

Genotyping mESC clones was performed either using PCR followed by gel electrophoresis as described or using real-time quantitative PCR (qPCR), which was performed with the KAPA SYBR FAST (Kapa Biosystems KK4610) on a LightCycler 480 Real-Time PCR System (Roche) using either 96-well or 384-well qPCR plates as previously described (Brosh et al. 2023). In most cases loading was performed using an Echo 550 liquid handler (Labcyte): a 384-well qPCR plate was prefilled with 5 µL KAPA SYBR FAST and 4 µL water per well. 100 nL of each 100 μM primer and 0.5 µL of each crude gDNA sample were transferred by the Echo. Genotyping payload clones typically included, in addition to assays designed to detect the newly formed left and right junctions, assays to detect the loss of the landing pad, the absence of the payload vector backbone, and for large (>100 kb) payloads, allele-specific assays to detect internal delivered regions. Genotyping primers are listed in **Table S6**.

### High-throughput sequencing verification

Illumina sequencing libraries were prepared from purified DNA as previously described (Brosh et al. 2023). The library construction method for each sample is listed in **Table S8**.

Hybridization capture (Capture-seq) for targeted resequencing of engineered regions was performed as previously described (Brosh et al. 2021). Briefly, biotinylated bait was generated using nick translation from BACs covering the *Igf2/H19* (RP23-50N22 or RP23-137N15), *Hprt* (RP23-412J16) or *Sox2* locus (RP23-144O8 or RP23-274P9) (see **Table S7** for BAC coordinates), landing pad plasmid (pLP-PIGA2), payload plasmid backbone, pCAG-iCre and pSpCas9 plasmids. Bait sets and sequencing statistics are listed in **Table S8**.

The sequencing parameters and the analysis pipeline are detailed in (Brosh et al. 2023). The full processing pipeline is also available at https://github.com/mauranolab/mapping.

### Payload sequence verification

Payload sequence verification to identify potential sample swaps or contamination was performed as previously described (Brosh et al. 2023). Briefly, mean sequencing coverage was calculated for the genomic regions corresponding to the payload sequence and to the non-payload sequence (i.e. the portion of the engineered locus not included in the payload). Mean coverage was then normalized by the payload vector backbone coverage. Coverage analysis results and quality control (QC) calls for each sample are reported in **Table S9** and summarized in **Table S2**.

To detect payload DNA variants that might have been introduced during assembly or propagation in yeast or bacteria, we analyzed variant calls relative to the payload custom reference having sequencing depth above 10 and quality score above 50 (DP ≥ 10 and QUAL ≥ 50). Payload variants are reported in **Table S10** and QC calls are summarized in **Table S2**.

### mESC clone sequence verification

mESC clone sequence verification was performed as previously described (Brosh et al. 2023) with slight modifications. Briefly, mean sequencing coverage depth was calculated for the genomic regions corresponding to the payload and non-payload (as defined above); and for the payload vector backbone, landing pad, landing pad backbone, and pCAG-iCre custom references. Coverage depth of engineered regions was normalized by the mean genomic coverage of their flanks (i.e. the un-engineered portion of the genomic region targeted by the bait), such as that a single copy corresponds to a value of ∼0.5. For landing pad clones, we expect normalized coverage depth of 0.5 ±0.3 for the non-payload region and <0.1 for landing pad backbone. For payload clones, the expected coverage depth was adjusted according to the expected ploidy of the site, such as the limits are ploidy / 2 ±0.3 for the payload region (0.2-0.8 for 1x; 0.7-1.3 for 2x; 1.2-1.8 for 3x), 0.5 ±0.3 for the non-payload region and <0.1 for the landing pad, payload vector backbone and pCAG-iCre. Coverage depth analysis QC calls for each clone are reported in **Table S11**.

We further verified the presence of BL6 allele variants, which are lost in regions replaced by landing pads and restored by delivered payloads, by calculating allelic ratios as the mean proportion of reads supporting the reference (BL6) allele (propREF) for the genomic regions described above. Expected proportions of the reference allele were adjusted according to ploidy to: 0.3-0.7 for 2x; 0.47-0.87 for 3x for payload regions, and 0-0.2 for non-payload regions. Regions with 10 or fewer variants were ignored. Allelic ratio QC calls for each clone are reported in **Table S11**.

To verify genomic integration sites, we detected sequencing read pairs mapping to two different reference genomes using *bamintersect* (Brosh et al. 2021) as previously described (Brosh et al. 2023) with slight modifications: same-strand reads mapping within 500 bp were clustered; a minimum width threshold of 125 bp was required for reporting; and junctions with few reads (ReadsPer10M < 20 or junctions representing less than 5% of the total ReadsPer10M) were excluded. *Bamintersect* junctions were classified hierarchically based on position (**Table S12**). Results and QC calls for each clone are reported in **Table S13**.

### Mesendodermal differentiation

Differentiation to mesendodermal cells was adapted from a published protocol (Thomson et al. 2011). mESCs were first recovered into 2i media for two days. On day 1 of differentiation, mESCs were seeded on plates coated with 0.1% gelatin (EMD Millipore ES006-B) in N2/B27 differentiation medium, containing a 1:1 mixture of DMEM/F12 (ThermoFisher 11330032) and Neurobasal (ThermoFisher 21103049), supplemented with 0.5% N2 Supplement (ThermoFisher 17502048), 1% B27 Supplement (ThermoFisher 17504044), 1% GlutaMAX (ThermoFisher 35050061), 1% Pen-Strep (ThermoFisher 15140122), 1% MEM nonessential amino acids (ThermoFisher 11140050), 100 μM sodium pyruvate (ThermoFisher 11360070), 0.1 mM 2-mercaptoethanol (Sigma M3148), supplemented with 10 μM ROCK inhibitor Y-27632 (Cell Signaling 13264). On day 2, ROCK inhibitor was removed and differentiation media was supplemented with 3 μM CHIR99021 (R&D Systems 4423) replenished daily for 3 additional days. On day 5, plates were rinsed twice using PBS and cells were collected for further studies.

### DNase-seq

DNase digestion was performed as described previously (John et al. 2013; Georgolopoulos et al. 2021) and adapted to 200 µL thermocycler tubes. Briefly, nuclei were extracted from cells and incubated with limiting concentrations of the DNA endonuclease DNase I (Sigma D4527-20KU) supplemented with Ca^2+^ and Mg^2+^ for 3 min at 37 °C. Digestion was stopped by the addition of EDTA, and the samples were treated with proteinase K and RNase A. Short double-hit fragments were isolated from DNaseI digestion using magnetic bead polyethylene glycol (PEG) fractionation. Illumina libraries were generated and sequenced on an Illumina NextSeq 500. Sequencing statistics are listed in **Table S8**.

### CUT&RUN

CUT&RUN experiments were conducted using the CUT&RUN (Cleavage Under Targets & Release Using Nuclease) assay kit (Cell Signaling Technology, #86652) following the manufacturer’s protocol. For each reaction, 5 × 10^6^ cells were used. Positive (H3K4me3) and negative (Rabbit mAb IgG) controls were performed using the antibodies provided in the kit. The CTCF antibody (Cell Signaling Technology, #2899S) was used at a dilution of 1/50 for overnight incubation at 4°C. DNA fragments were purified using the DNA Clean & Concentrator-5 Kit (Zymo, D4014) according to the manufacturer’s procedures, and eluted in 15 µL. Up to 11.15 µL purified DNA was taken directly into end repair for library construction using the Illumina dsDNA protocol as previously described (Brosh et al. 2023). During PCR enrichment of adaptor-ligated DNA, the extension time was reduced to 15 sec to exclude amplification of large library DNA fragments (>1,000 bp) and reduce background signal. The sequencing parameters and the analysis pipeline are detailed in (Halow et al. 2021). The full processing pipeline is also available at https://github.com/mauranolab/mapping. Sequencing statistics are listed in **Table S8**.

#### RNA Isolation, cDNA synthesis and mRNA expression analysis by real-time qRT-PCR

Expression assays were performed as previously described (Brosh et al. 2023). Briefly, RNA was isolated using the RNeasy-mini (QIAGEN 74106) protocol followed by DNase treatment using the Turbo DNA-free kit (ThermoFisher AM1907). cDNA was synthesized from 1-2 μg total RNA with the Multiscribe High-Capacity cDNA Reverse Transcription Kit (ThermoFisher 4368814).

Real-time quantitative reverse transcription PCR (qRT-PCR) was performed using KAPA SYBR FAST (Kapa Biosystems KK4610) on a 384-well LightCycler 480 Real-Time PCR System (Roche) and threshold cycle (Ct, also called Cp) values were calculated using Abs Quant/2nd Derivative Max analysis. Primers were designed to detect *Igf2* and *H19* transcripts in an allele-specific manner (**Figure S6** and **Table S6**). In most cases loading was performed using an Echo 550 liquid handler (Labcyte): a 384-well qPCR plate was prefilled with 5 µL KAPA SYBR FAST and 4 µL water per well. 100 nL of each 100 μM primer and 1 µL of each cDNA sample were transferred by the Echo. Thermal cycling parameters were as follows: 3 min pre-incubation at 95 °C, followed by 40 amplification cycles of 3 sec at 95 °C, 10 sec at 60 °C and 10 sec at 72 °C. *Sox2* expression was quantified using a similar protocol but with annealing temperature lowered to 57 °C as in (Brosh et al. 2021).

Assays were performed in duplicate and replicate wells on the same plate were averaged after masking wells with no or very low amplification (Cp > 30) and low RNA quality (Gapdh Cp > 19). Raw Ct values are listed in **Data S2**. ΔCt values for the BL6 and CAST alleles were computed relative to *Gapdh* (*Hprt* in the time course differentiation experiment). Clones demonstrating nonspecific differentiation under ESC culture conditions were identified as having high *Igf2* CAST allele expression (ΔCt < 7 in ESC) and discarded (n=5). Replicate ΔCt measurements of the same clone across different plates and differentiations were averaged. Activity was reported as 2^-ΔCt^, scaled to yield expression values ranging from 0 (Δ*Igf2/H19*) and 1 (“Full-length”) (**Table S5**). Scaling was performed by subtracting the median of the Δ*Igf2/H19* samples in 2i, then dividing by the difference between the median of the appropriate “Full-length” payload sample: constitutive CTCF occupancy for *H19,* and abolished CTCF occupancy for *Igf2* and the median of the Δ*Igf2/H19* samples.

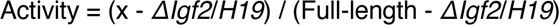

Sox2 expression was scaled similarly, and as previously described in (Brosh et al. 2023). Briefly, *Sox2* fold change was defined as the difference between the BL6 and CAST *Sox2* alleles, calculated as 2^ΔCt[CAST-BL6]^. Fold change was scaled to yield expression values ranging from 0 (ΔSox2) to 1 (“*Sox2* & Sox2LCR”) by subtracting the mean fold change calculated for the ΔSox2 samples from all data points, then dividing by the mean fold change of the appropriate wild-type payload samples (“*Sox2* & Sox2LCR” or “Sox2LCR”).

#### Transcriptional analysis of mesendoderm differentiated cells

Transcriptomic analysis of mesendoderm differentiated cells was performed in 6 independent engineered clones: 3 clones including the “Full-length” payload with constitutive CTCF occupancy at *Igf2/H19* and 3 clones including “Full-length” payload with abolished CTCF occupancy. RNA-seq libraries were prepared from 500 ng of total RNA using the NEBNext Ultra II Directional RNA Library Prep Kit for Illumina (NEB E7760S), and performing PolyA selection for ribosomal depletion using NEBNext Poly(A) mRNA Magnetic Isolation Module (NEB E7490) according to the manufacturer’s protocol. Libraries were sequenced in single-end mode on a NextSeq 500 (**Table S8**).

Transcriptional profiles from 6 mESC replicates were analyzed from published RNA-seq data. Reads were aligned to the mouse genome build mm10 using STAR and mapped to transcripts using RSEM. Raw reads counts were normalized using DESeq2 and differential expressed genes were determined as p < 0.05, |FC| > 1. Enrichment analysis of protein coding genes was done using clusterProfiler on published sets of genes with high or low expression in differentiated mesendoderm (p < 0.05) (Shen et al. 2008).

#### Cross-cell type DHS correlation analysis

Tissue-specific accessibility patterns were based on a master peak list of 45 representative samples from mouse ENCODE DNase-seq data (Halow et al. 2021). The cell-type activity spectrum (MCV, for multi-cell verified) was computed using ‘bedmap --count’ to consider overlaps with the center ±25 bp of each feature. MCV was then scaled from 0 to 1 by dividing by 45.

Accessibility per DHS peak per cell/tissue type was quantified using bedmap --max from density tracks. The Pearson correlation coefficient (r) was calculated pairwise between each viewpoint and each DHS within the surrounding 1-Mb window. We then fitted a loess regression curve (span = 0.2). The coordinates for the *Sox2* LCR correspond to the DHS23 to DHS26 (Brosh et al. 2023) (**Table S7)**. Gene TSS viewpoints were computed as TSS ±1 kb.

DHS accessibility correlation was assessed for three viewpoint sets: (i) 1000 random DHSs selected from a peak master list of 45 representative samples from mouse ENCODE DNase-seq data (Halow et al. 2021) and located at least 10 kb away from a gene TSS; (ii) promoters of protein-coding genes (±1 kb from TSS, Gencode release M23, genes located in mitochondrial DNA were excluded); (iii) mESC super-enhancers (Whyte et al. 2013). Viewpoints with less than 80% mappable sequence in the 1 Mb surrounding window were excluded. DHS correlation patterns were grouped based on the number of instances within a 1-Mb window that the smoothed Pearson correlation exceeded 0.25: 0 times (uncorrelated), 1 time (independent), and 2+ times (coordinated).

## Supplementary Materials

### SUPPLEMENTARY FIGURES

**Figure S1.**
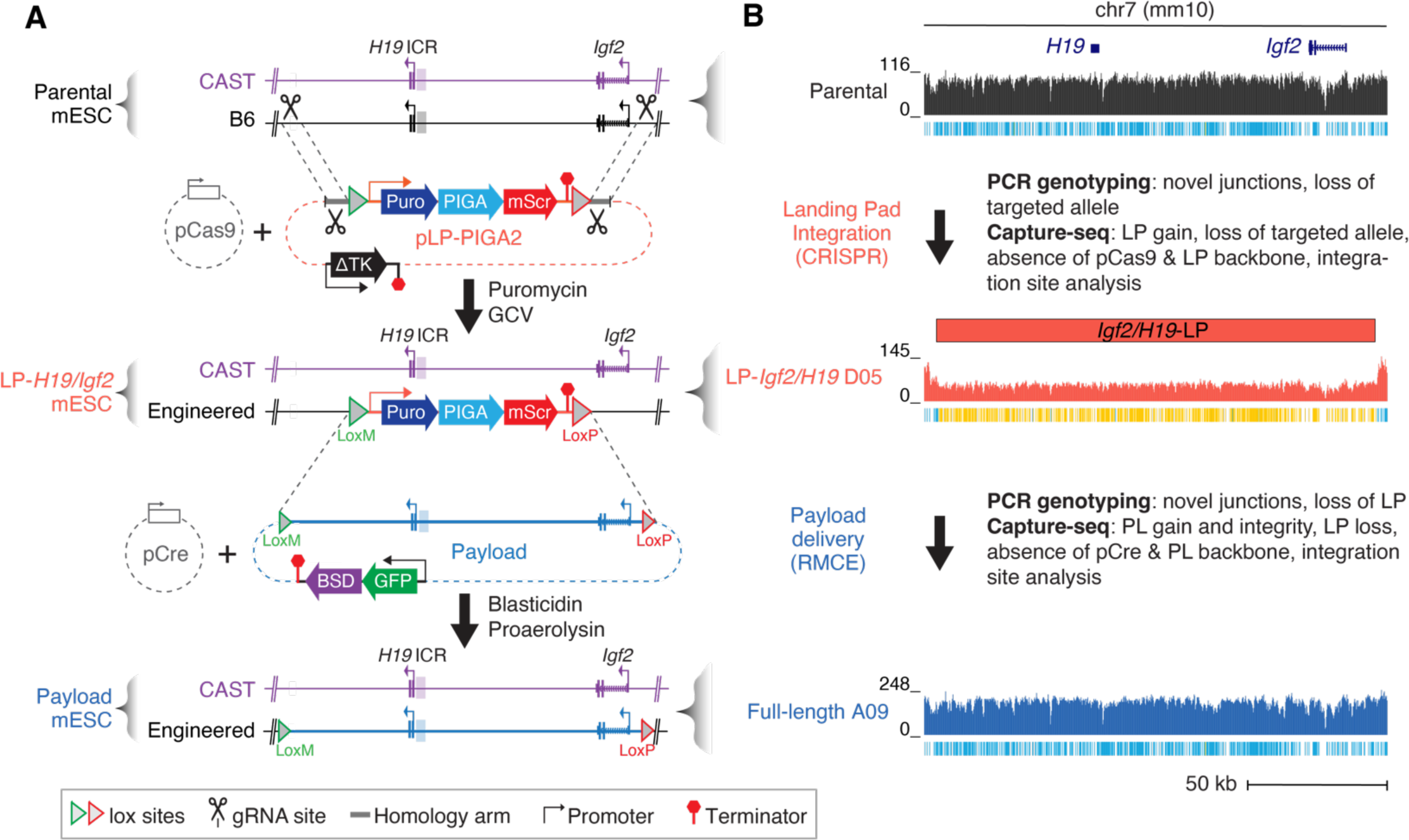
Big-IN landing pad at *Igf2/H19*. (**A**) Engineering strategy for LP-*Igf2/H19* mESCs. The B6 allele of the 157-kb *Igf2/H19* locus was replaced with a landing pad (LP-PIGA2). Landing pad integration was driven by Cas9 using a pair of gRNAs targeting both the replaced allele and the landing pad plasmid and by short homology arms that facilitate homology-directed repair. Landing pad mESCs were selected with puromycin, while ganciclovir (GCV) selects against landing pad backbone integration. Cre recombinase-mediated cassette exchange (RMCE) enabled replacement of the landing pad with a series of payloads. Payload-transfected cells were transiently selected with blasticidin, followed by counterselection of remaining landing pad mESCs cells with proaerolysin. (**B**) Schematic of mESC clone verification pipeline. Representative examples of targeted capture sequencing validation throughout the engineering process: parental (ΔPiga) mESCs, LP-*Igf2/H19* clone D05, and full-length A09 payload clones. Ticks under each coverage track indicate heterozygous (B6xCAST, blue) or homozygous non-reference (CAST, yellow) variants.

**Figure S2.**
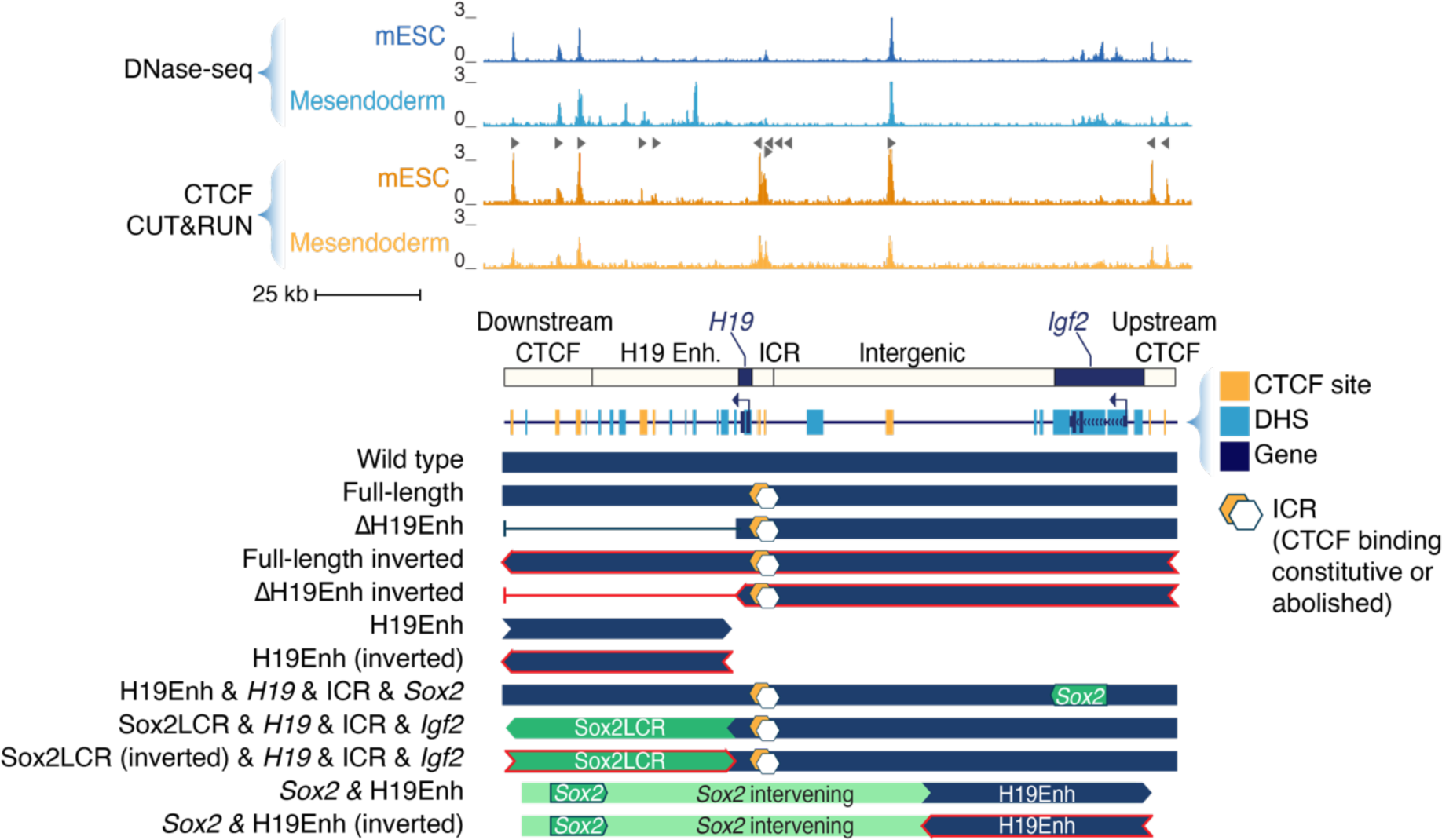
Schematic representation of engineered payloads. Shown are DNase-seq and CTCF CUT&RUN tracks and the CTCF sites and DNase Hypersensitive Sites (DHS) identified throughout the *Igf2/H19* locus. Inverted sequences are colored in red. Hexagons denote payloads pairs that are identical except for their ICR structure.

**Figure S3.**
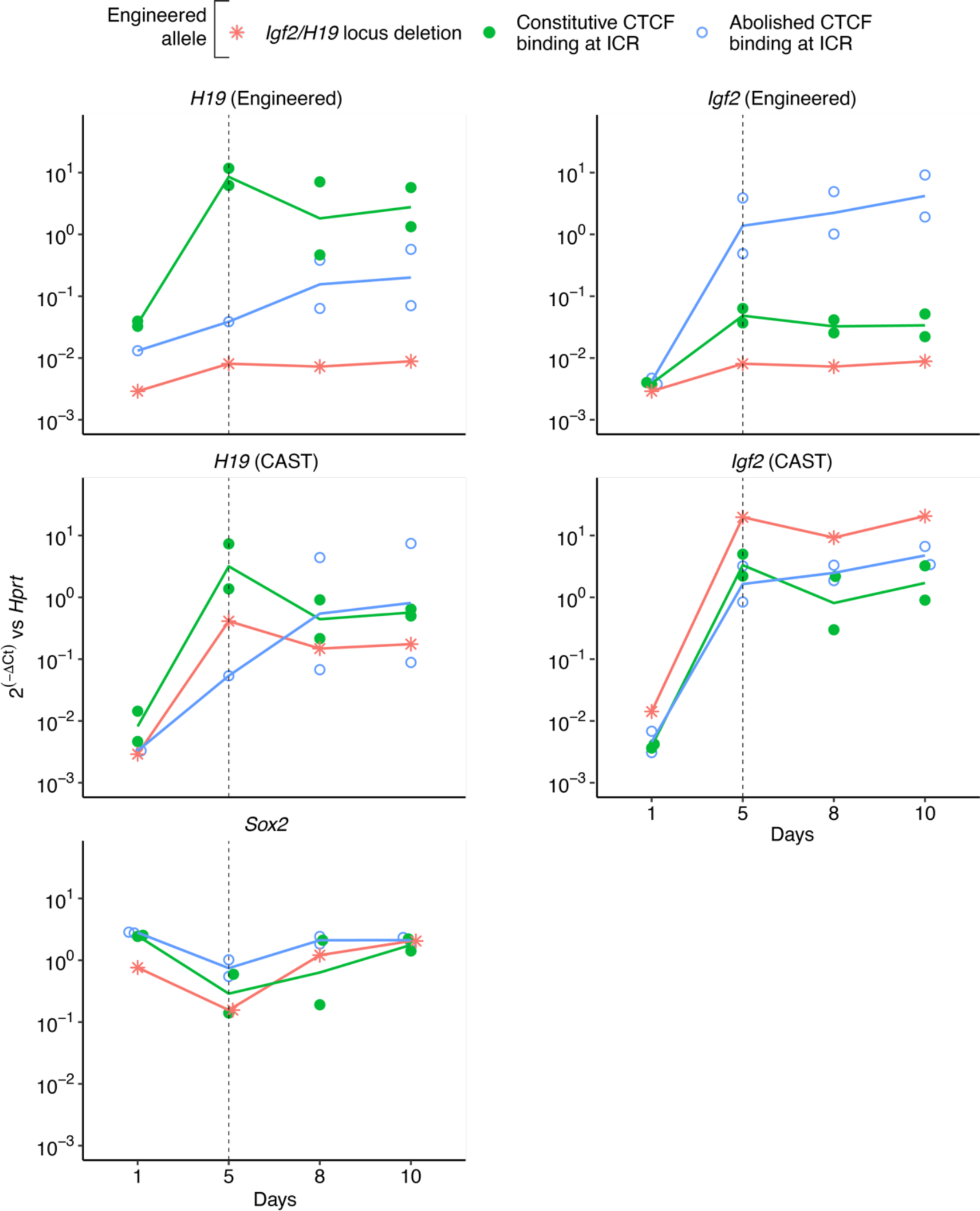
Mesendodermal differentiation time course. Expression analysis using qRT-PCR during a differentiation time course from mESC to mesendodermal cells. *H19* and *Igf2* expression was measured using allele-specific primers for the engineered and the CAST alleles, and *Sox2* expression using primers that amplify both alleles. Each point represents the expression of an independently engineered mESC clone normalized to *Hprt*. Lines represent medians. The selected time point for further differentiation experiments is represented as a vertical dashed line.

**Figure S4.**
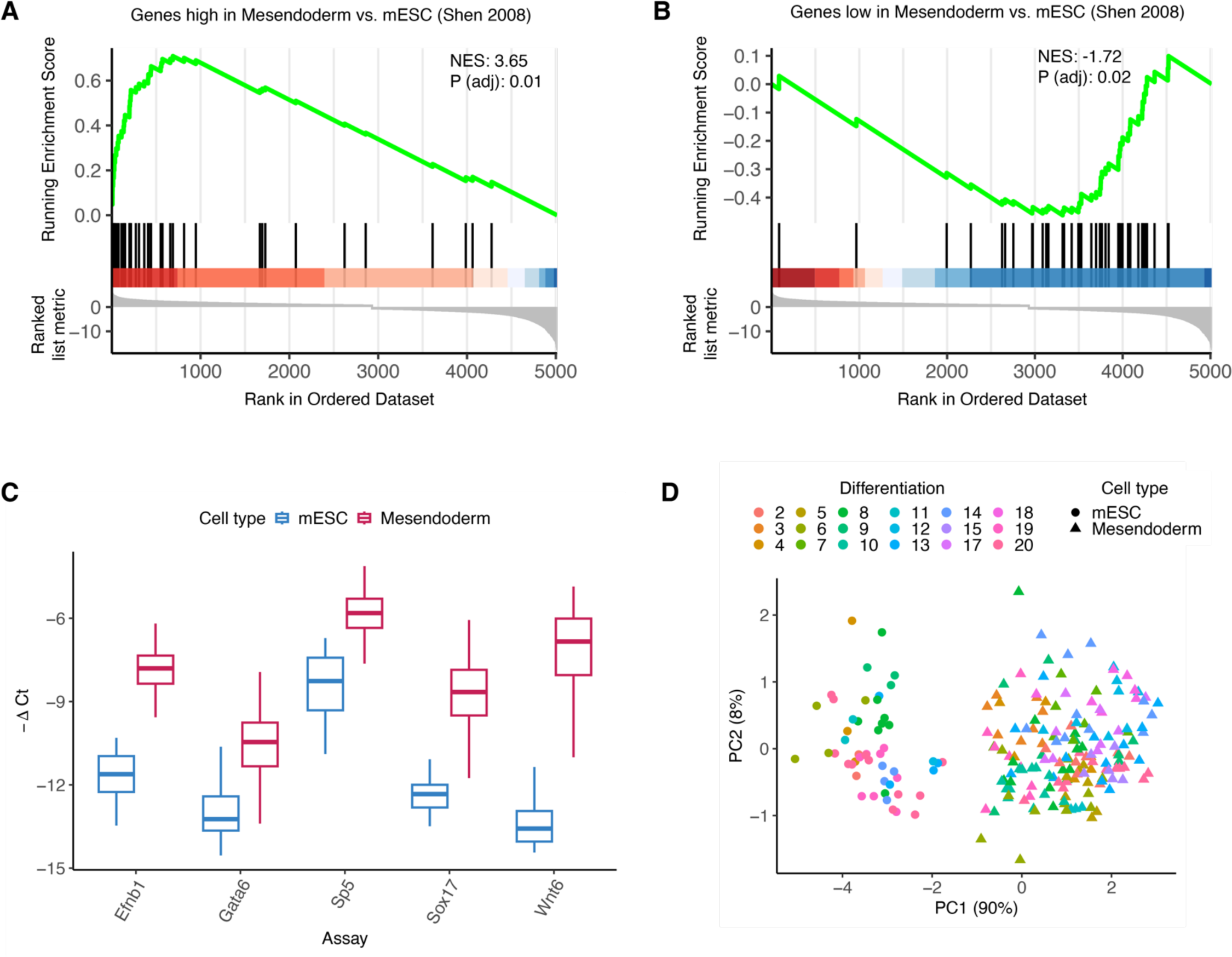
Mesendoderm marker gene expression profiling. (**A**-**B**) Gene set enrichment analysis of differentially expressed genes in RNA-seq data from mesendoderm differentiated cells vs. mESC. The two sets of genes high (**A**) and low (**B**) in mesendoderm were obtained from gene expression profiles from (Shen et al. 2008). NES, Normalized Enriched Score. (**C**-**D**) qRT-PCR expression analysis of mesoderm marker genes identified from RNA-seq. (**C**) Quantification across all clones in mESC and differentiated cells. (**D**) Principal component analysis. Clones are colored by differentiation batch, and shape denotes mESC (circles) and mesendodermal cells (triangles).

**Figure S5.**
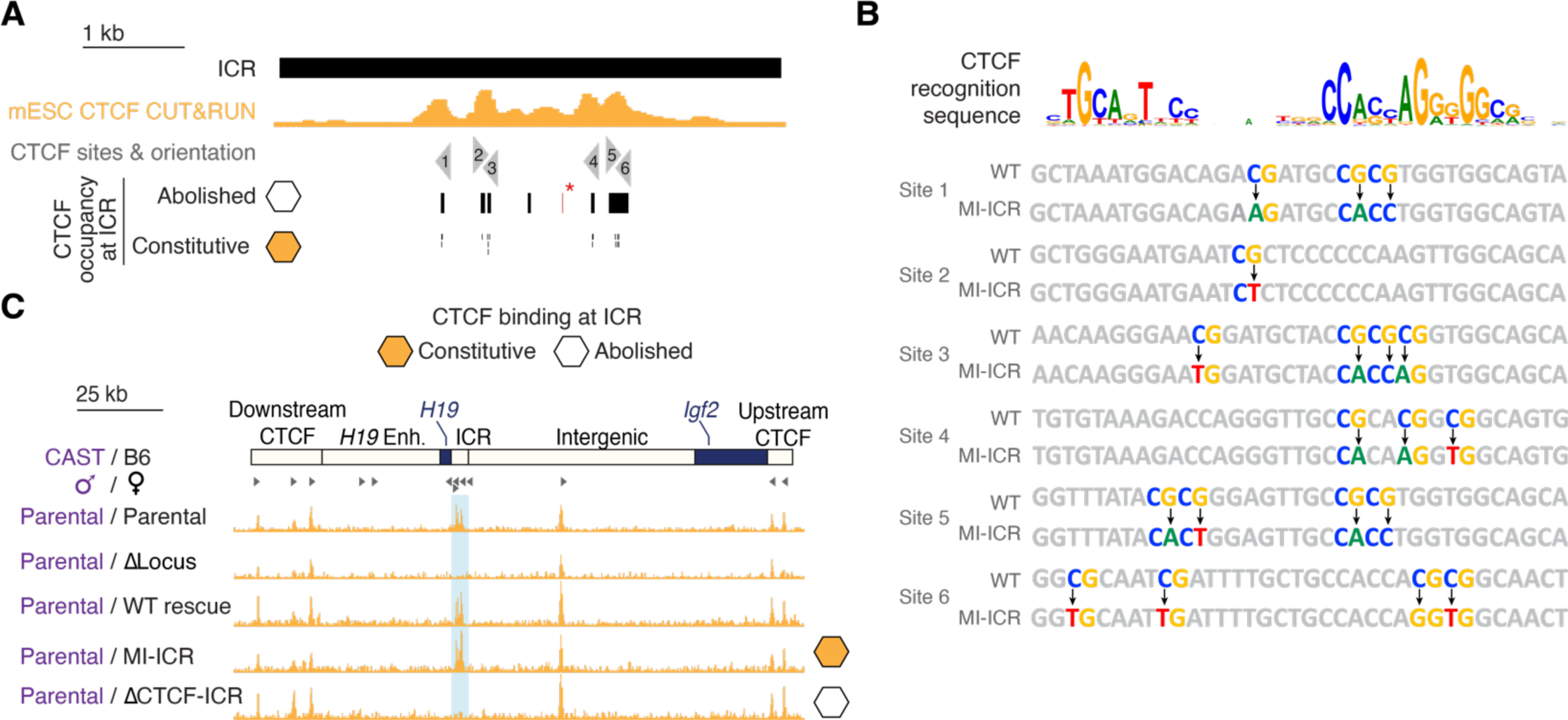
Engineering the imprinting control region to control CTCF occupancy. (**A**) Detail of the *Igf2/H19* ICR showing CTCF occupancy in mESC. Triangles indicate CTCF recognition sequence orientation. Assembled payloads included either CTCF site deletions resulting in abolished CTCF occupancy (ΔCTCF-ICR, empty hexagon) or CpG mutations resulting in constitutive CTCF occupancy (methylation-insensitive ICR or MI-ICR, orange hexagon). *, Point deletion detected in the “Full-length” ΔCTCF-ICR payload (see BS09866A in **Table S10**); this point variant is not expected to confound the measured activities of the integrated payload. (**B**) Detail on the methylation-insensitive mutations in the MI-ICR design where all CpGs in the 6 CTCF recognition sequences were mutated without significant alteration of the CTCF canonical recognition sequence. (**C**) CUT&RUN CTCF signal at the *Igf2/H19* locus in differentiated mesendodermal cells. Genotypes for each track are indicated at the left. The shaded region denotes the ICR. The CTCF recognition site orientations are indicated at top by triangles.

**Figure S6.**
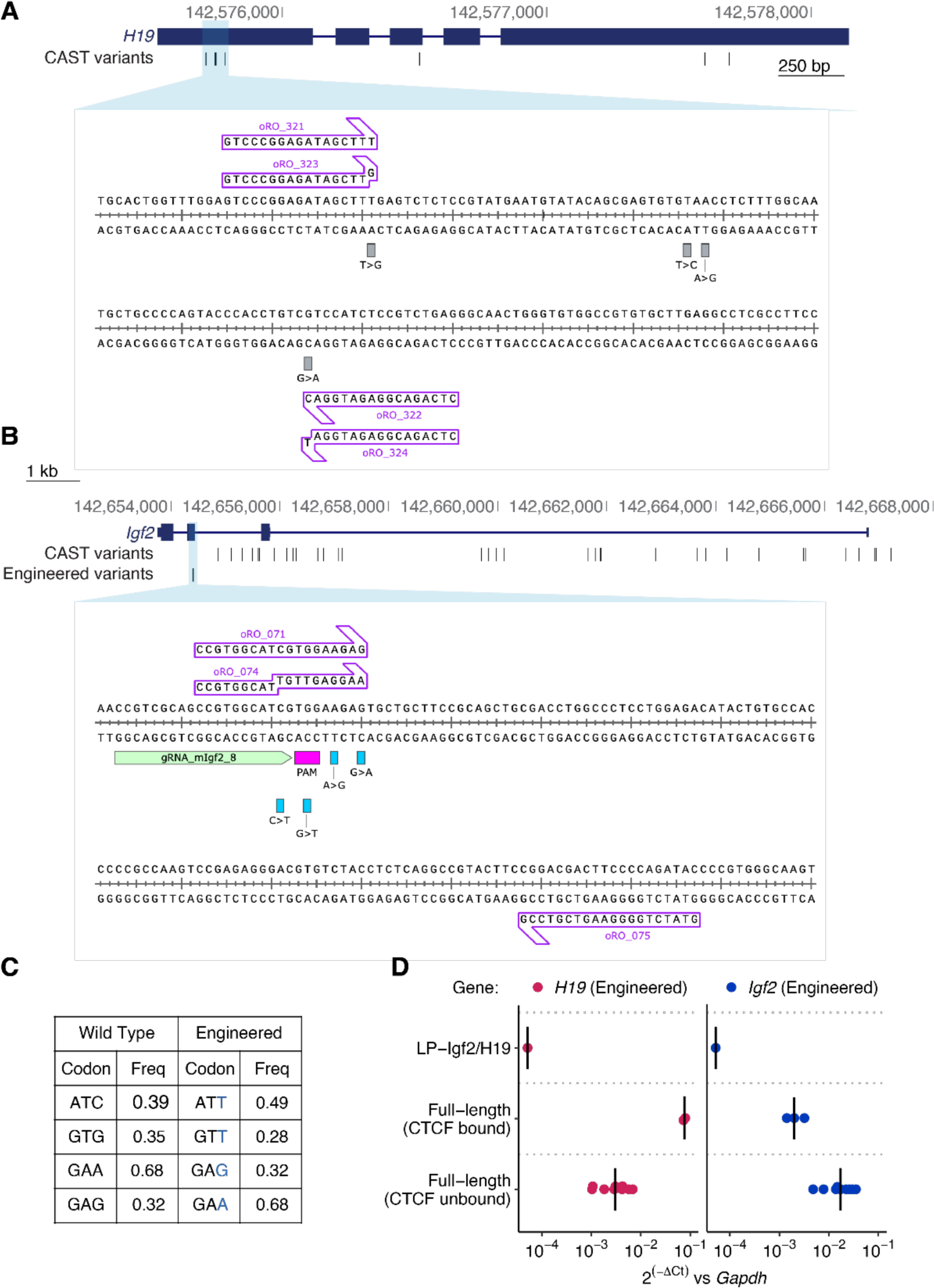
Allele-specific qRT-PCR expression analysis. (**A**) Schematic representation of the *H19* gene, including the CAST variants present in the unengineered allele and a detailed sequence view including the allele-specific primers used in the qRT-PCR assays (as detailed in **Table S6**). (**B**) Schematic representation of the *Igf2* gene. Shown below are CAST variants present in the unedited allele, and engineered synonymous variants in exon 4. The detailed sequence view includes the allelespecific primers used in the qRT-PCR assays (as detailed in **Table S6**) and the gRNA used for payload engineering. (**C**) Codons overlapping the 4 point substitutions in the wild-type and engineered *Igf2* showing their genomic frequencies. (**D**) Allele-specific qRT-PCR assay for *H19* and *Igf2* expression in differentiated mesendodermal cells. Transcripts from engineered alleles were measured using allele-specific primers in LP-*Igf2*/*H19* as well as “Full-length” payload clones with constitutive or abolished CTCF binding to the ICR. Each point represents the expression of the engineered *H19* or *Igf2* gene in an independent clone, measured using allele-specific qRT-PCR. Bars indicate median.

**Figure S7.**
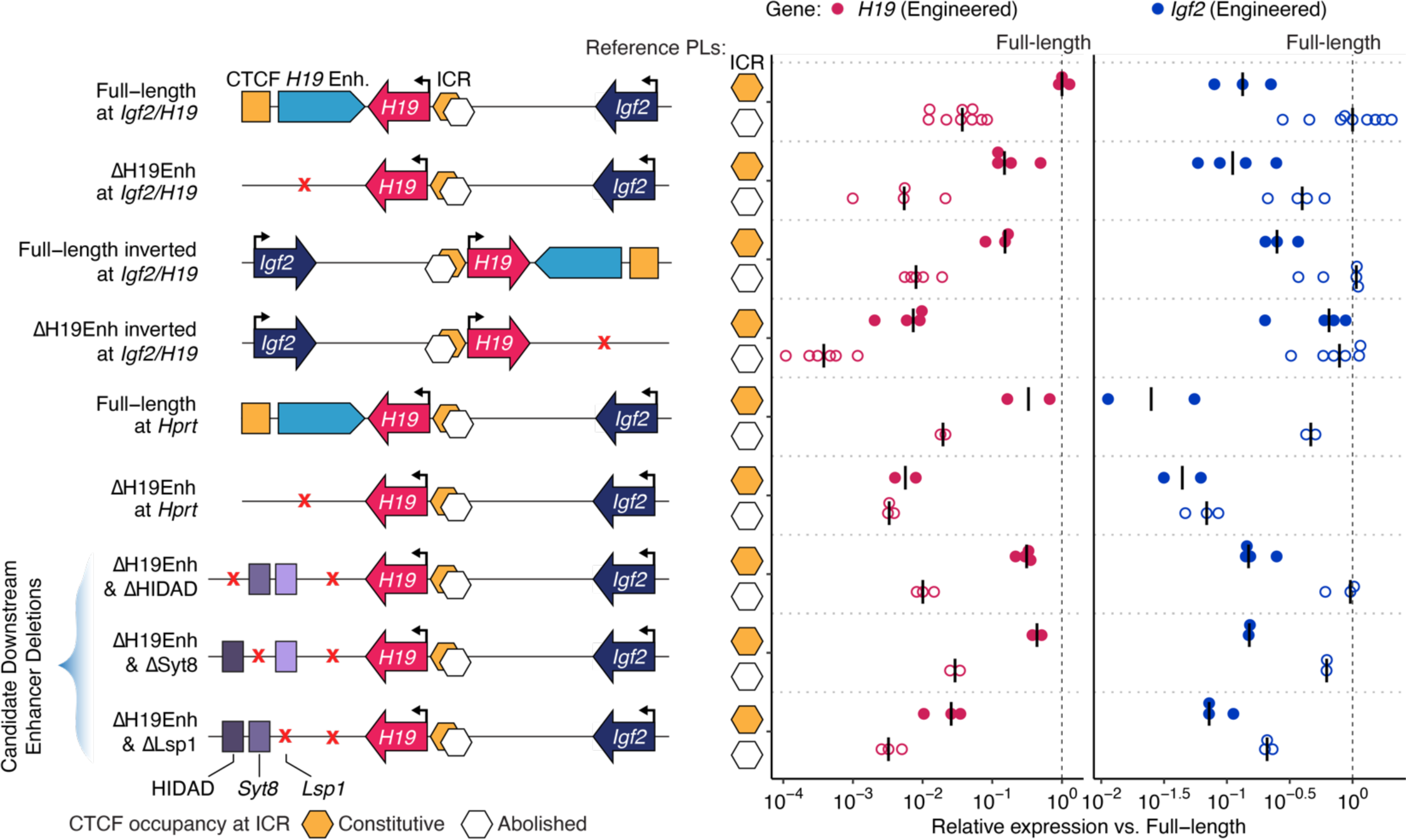
*Igf2/H19* payloads expression analysis on logarithmic scale. Expression was normalized to *Gapdh* and then to the reference “Full-length” rescue payloads, which are also represented as dashed lines (“Full-length” payload with constitutive CTCF occupancy for *H19* and “Full-length” payload with abolished CTCF occupancy for *Igf2*). Each point represents the expression of the engineered *H19* or *Igf2* in an independent clone using allele-specific qRT-PCR with bars indicating the median. Payloads with both constitutive CTCF occupancy (orange hexagon) or abolished CTCF occupancy (empty hexagon) at the ICR were assessed. Data are the same as in **Figure 1E**, **Figure 2B**, and **Figure S6D**.

**Figure S8.**
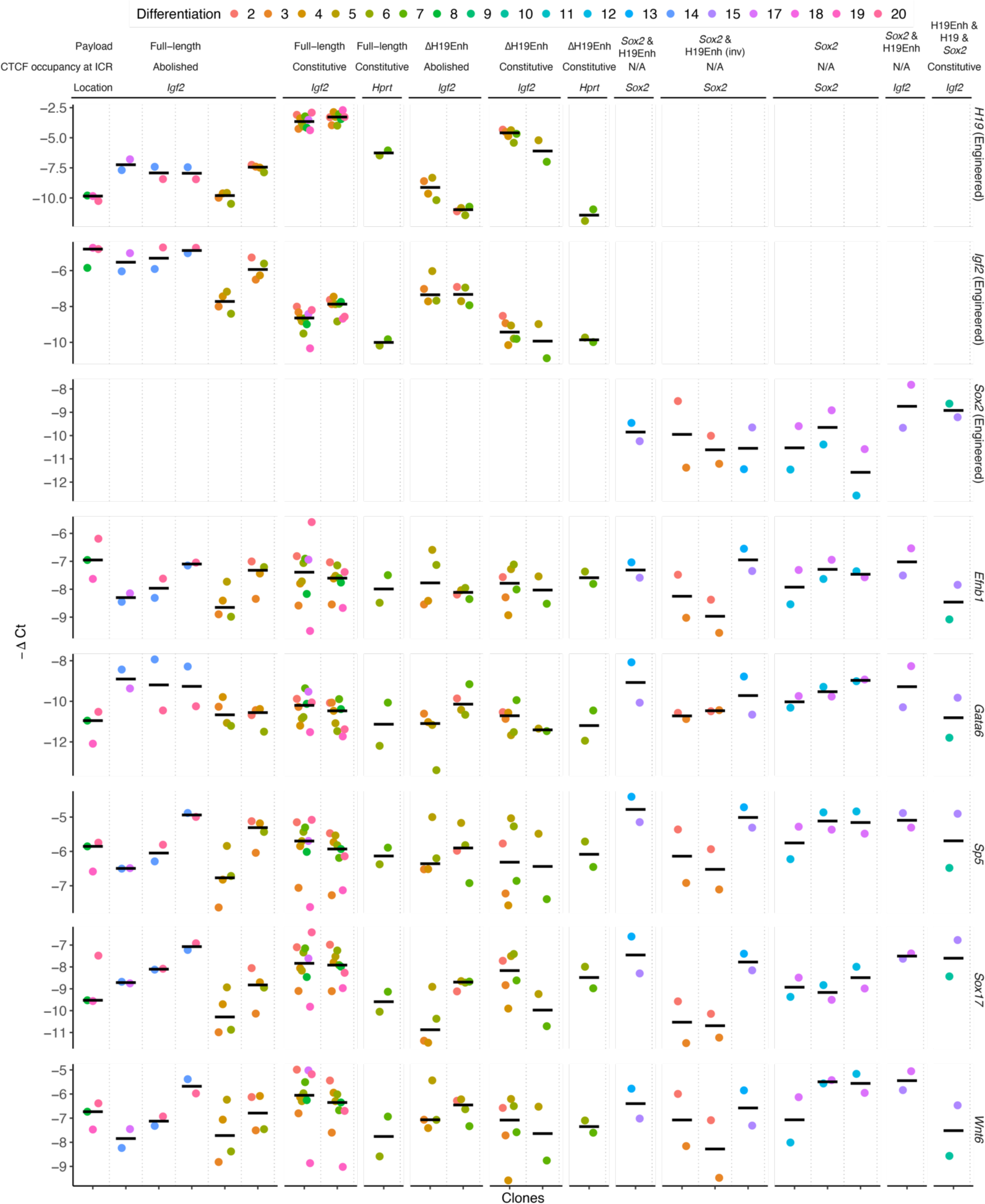
Expression consistency analysis in clones differentiated multiple times. Shown are mesendoderm marker gene and engineered gene (*H19*. *Igf2*, and *Sox2*) expression by differentiation batch. Independent clones are listed along the x-axis, and each point represents a unique differentiation batch. Payload ID and delivery location are listed above.

**Figure S9.**
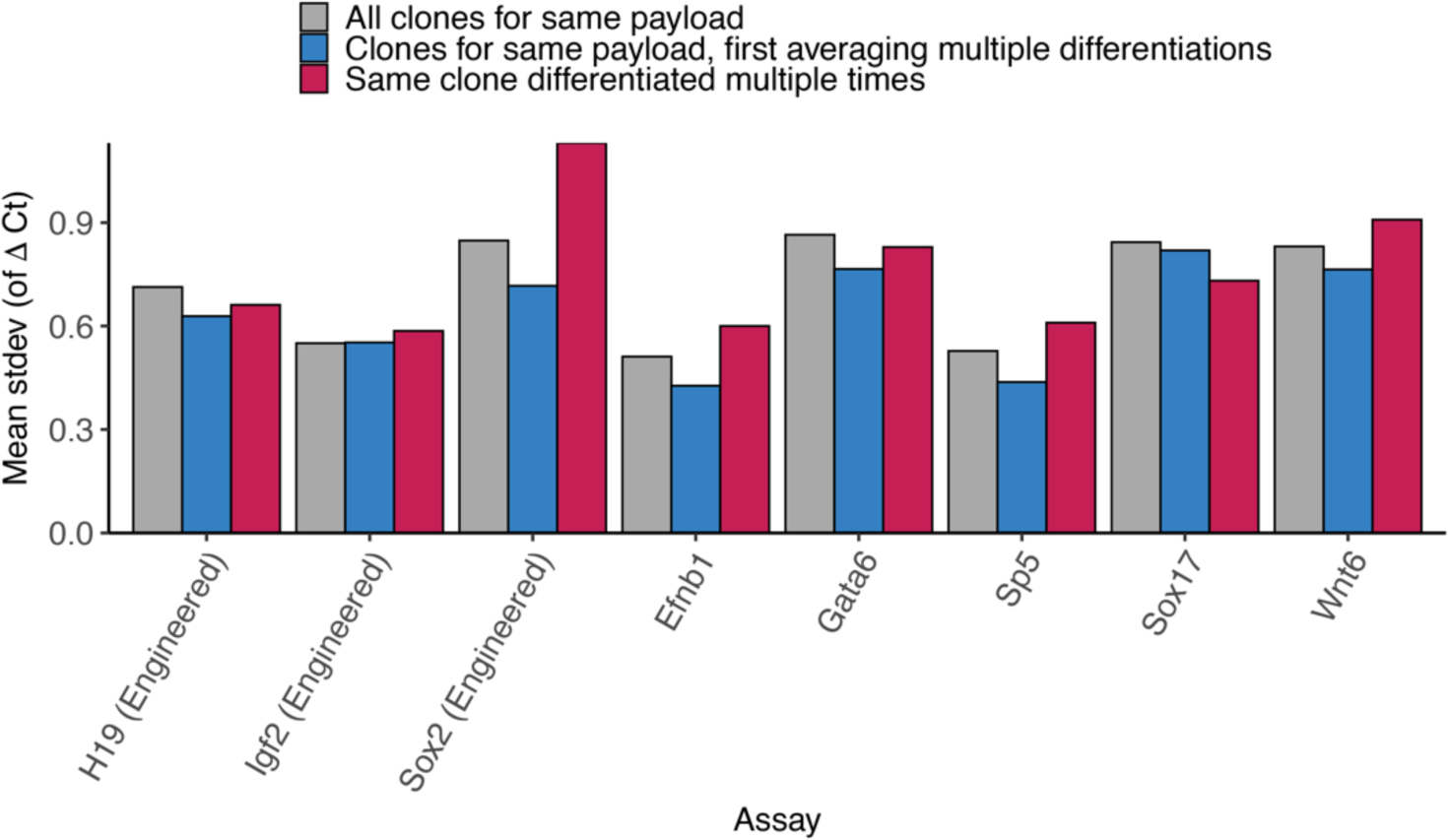
Replicate variability analysis. Analysis of standard deviation (stdev) of mesendoderm marker and engineered gene expression across replicates. Replicates were considered as multiple clones for the same payload (gray), multiple clones for the same payload while first averaging data from different differentiations (blue), or across differentiations for the same clone (red). Variation across differentiations was computed only for clones differentiated multiple times.

**Figure S10.**
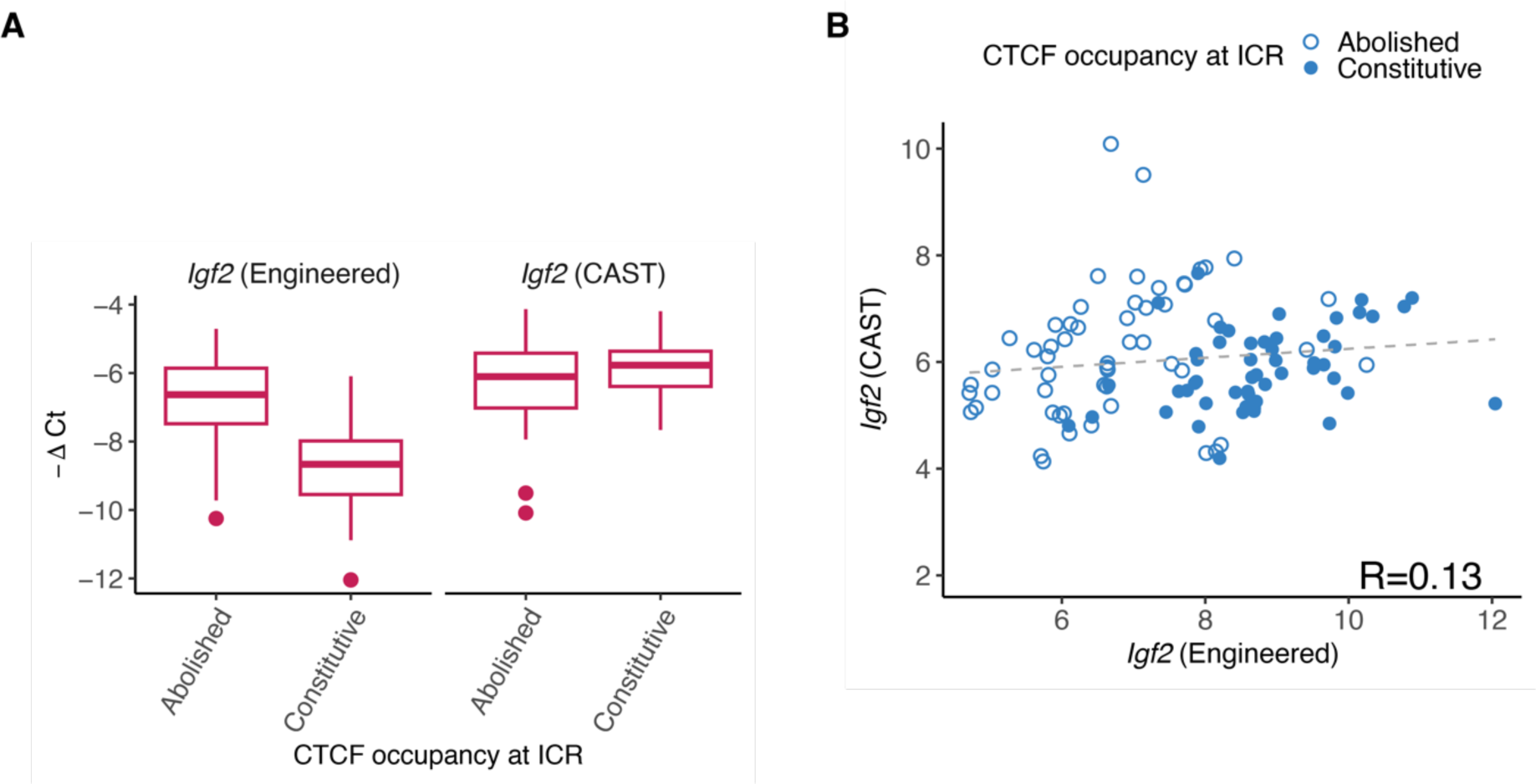
Expression of engineered versus CAST allele of *Igf2* in mesendoderm. (**A**) Expression of all payloads, grouped by whether the payload contained an ICR with abolished or constitutive CTCF occupancy (x-axis). CTCF occupancy at the ICR reduced expression of the Engineered but not CAST *Igf2* allele. (**B**) Analysis of the correlation between Engineered and CAST alleles in the same clone. Each point represents a distinct clone.

**Figure S11.**
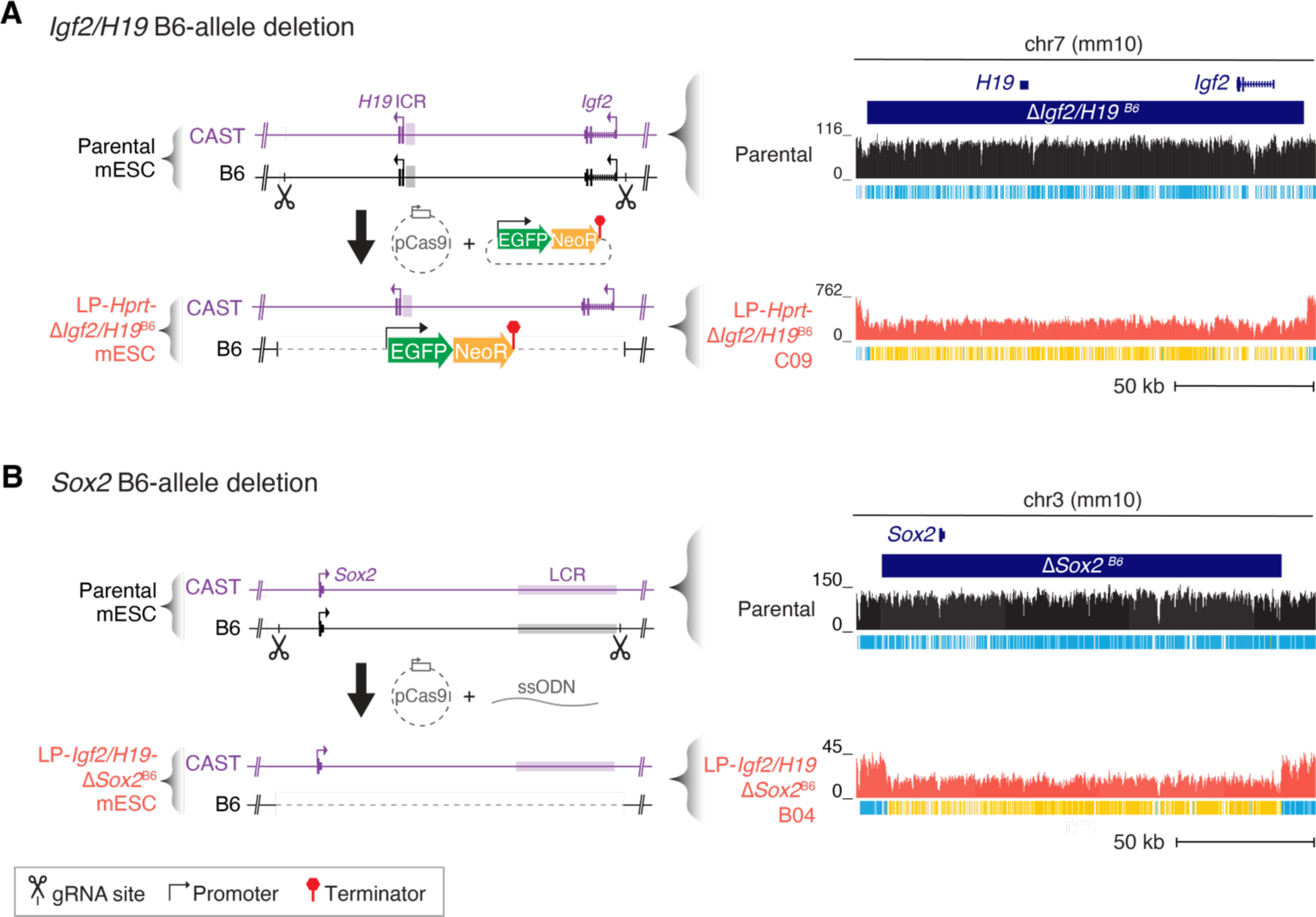
*Igf2/H19* and *Sox2* deletions. (**A**) Engineering of *Igf2/H19* B6-allele deletion. The B6 allele of the *Igf2/H19* locus in chromosome 7 was deleted through replacement by pDONOR∼Neo∼EGFP. Representative examples of targeted capture sequencing validation are shown: coverage tracks for parental (ΔPiga) mESCs and LP-*Hprt*-Δ*Igf2/H19* mESCs including deletion of the B6 *Igf2/H19* allele (Δ*Igf2/H19*^B6^) clone D05. Ticks under each coverage track indicate heterozygous (B6xCAST, blue) or homozygous non-reference (CAST, yellow) variants. (**B**) Engineering of *Sox2* B6-allele deletion. The B6 allele of the *Sox2* locus at chromosome 3 was deleted using Cas9 and a single-stranded oligodeoxynucleotide (ssODN) as a donor for homology-directed repair. Representative examples of targeted capture sequencing validation are shown: coverage tracks for parental (ΔPiga) mESCs and LP-*Igf2/H19*-Δ*Sox2* mESCs including deletion of the *Sox2* B6 allele clone B04. Ticks under each coverage track indicate heterozygous (B6xCAST, blue) or homozygous non-reference (CAST, yellow) variants. Puro, puromycin-resistance gene; PIGA, human mini *PIGA* gene; mScr, mScarlet; ΔTK, truncated thymidine kinase; BSD, blasticidin-resistance gene; eGFP, enhanced green fluorescent protein gene; NeoR, Neomycin-resistance gene.

**Figure S12.**
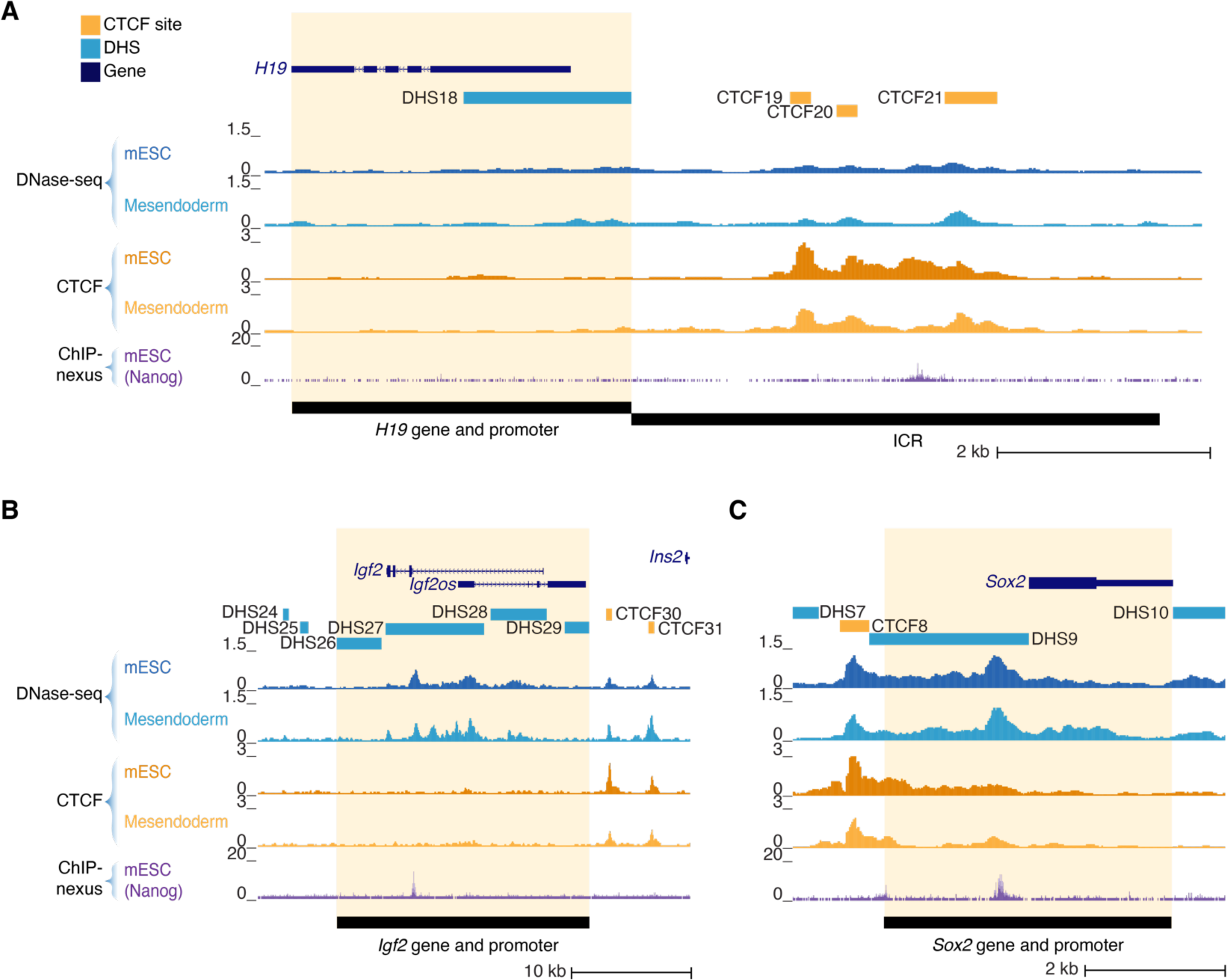
*H19*, *Igf2* and *Sox2* gene and promoter regions. Genomic region surrounding *H19* gene and the ICR (**A**), *Igf2* gene (**B**), and *Sox2* gene (**C**), showing DNase-seq and CTCF CUT&RUN in mESC and differentiated mesendodermal cells, together with ChIP-nexus data for Nanog transcription factor occupancy in mESC. The region engineered in those payloads including gene and promoter swaps is represented by the black bar and highlighted in yellow (see **Table S7** for region coordinates).

**Figure S13.**
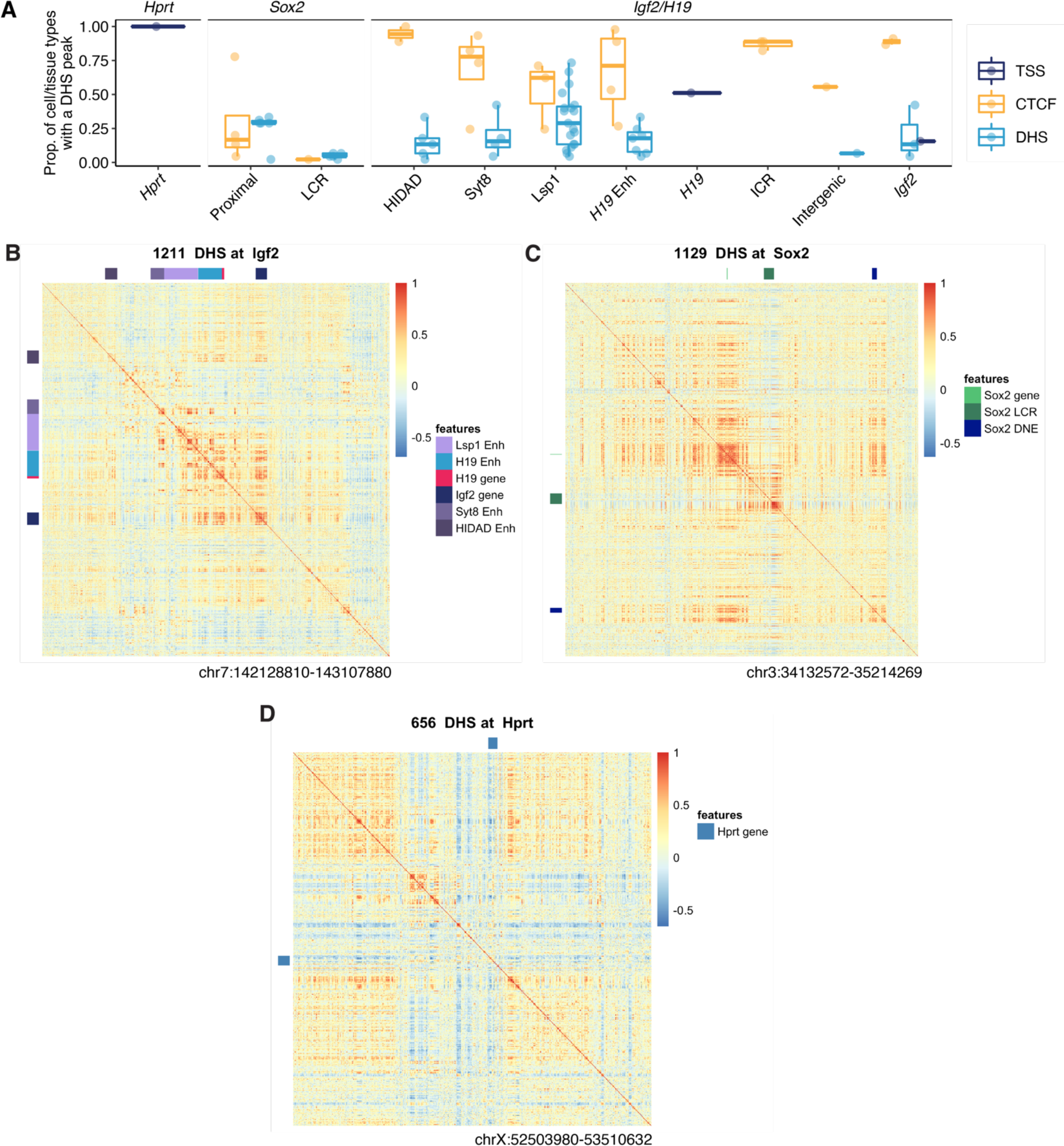
DHS accessibility correlation. (**A**) Cell and tissue selectivity of DHSs at the *Hprt*, *Sox2*, and *Igf2*/*H19* loci. Accessibility was assessed as the proportion of 45 samples from mouse ENCODE which had a DHS. DHSs overlapping CTCF sites and TSS were analyzed separately. Bars represent the median, boxes represent the interquartile range, and whiskers the minimum and maximum values. (**B-D**) Heatmap of pairwise correlations (Pearson r coefficient) of DHS accessibility in the 1-Mb regions surrounding *Igf2/H19* (**B**), *Sox2* (**C**) and *Hprt* (**D**). Correlations were computed across 45 mice ENCODE DNase-seq samples. Relevant regulatory elements or genes are labeled with distinct colors. DNE, Distal Neural Enhancer (Chakraborty et al. 2023).

## SUPPLEMENTARY TABLES

**Table S1. Payloads and assembly details.**

Strategies and reagents used for payload, as well as sequencing data. See **Methods** for details of each assembly strategy, **Table S6** for cloning primer sequences, gRNA sequences and synthetic fragment sequences. Source or intermediate payloads not delivered to mESCs were assigned “Intermediate” in the “Assembly Step” field and those not including an ICR sequence in the design are assigned n/a in the “ICR design” field. Payload ID, a unique name for each payload.

This table is provided as a separate file.

**Table S2.** Summary of payload sequencing QC.

| Delivered payloads (n=19) |  |  |  |
| --- | --- | --- | --- |
|  | <u>Passed</u> | <u>Failed</u> | <u>No call</u> |
| Coverage | 18 (94.7%) | 1 (5.3%)(a) | 0 |
| Variant calls | 19 (100%) | 0 | 0 |

**Table S3. mESC clones.**

Summary of mESC clones described in this manuscript. Genetic Modification details DNA elements integrated and their integration site in square brackets; transiently transfected plasmids (e.g. pSpCas9 or pCAG-Cre) are indicated by null. See **Table S5** for payload labels used in figures. “Bl6/C_LP300(Igf2_1_4)_D5” corresponds to LP-*Igf2/H19* mESCs, “BL6/C LP300(Igf2_1_4)_D5 dSox2_201026_B4” corresponds to LP-*Igf2/H19-*Δ*Sox2* mESCs including the deletion of the *Sox2* B6 allele, “BL6/C LP300(Hprt_3g_4g)_20210414_C04” corresponds to LP-*Hprt* mESCs, “Bl6/C_LP300(Hprt_3_4)_C04_dIgf2_C9” corresponds to LP-*Hprt-*Δ*Igf2/H19* cells including deletion of the *Igf2/H19* B6 allele, “BL6/C LP300(Sox2_1g_5g)_201007_D4” corresponds to LP-*Sox2* mESCs, and “Bl6/C LP305(Sox2_6g_7g)_20210326_C01” corresponds to LP-LCR mESCs. Sample ID, a unique sequencing library identifier; Clone ID, a unique descriptor of the mESC clone; Payload ID, LIMS name for relevant payload or landing pad; Source, publication in which this mESC clone was first described.

This Table is provided as a separate file.

**Table S4. Summary of RNA-seq sequencing libraries.**

Sample ID, a unique sequencing library ID; Repository ID, a unique ID RNA-seq data obtained from the GEO repository; Clone ID, a unique descriptor of mESC clones; Source, unique identifier for a specific differentiation experiment date. For each sample, the Number of pass-filter alignments, Uniquely mapped and Nonredundant reads is provided. Batch indicates the differentiation batch.

This Table is provided as a separate file.

**Table S5. qPCR expression values and activity.**

*Igf2, H19* and *Sox2* expression in engineered clones. Payload ID, a unique name for each payload. Location, delivery location in the genome. Additional Deletions, presence of additional deletions engineered in that mESC clone. Label, display name used in Figure panels. Clone ID, a unique identifier for each engineered mESC clone. Cell Type, indicates whether expression was measured in mESC or mesendodermal cells differentiated for 5 days (Mesendoderm_5). Assay, targeted gene. deltaCt, ΔCt values relative to *Gapdh* for each gene. Activity, expression quantification as detailed in **Methods**.

This Table is provided as a separate file.

**Table S6. Primers, gRNAs, and synthetic DNA fragments.**

Primer pairs (qRT-PCR, genotyping PCR, amplification of DNA fragments for payload assembly, homology arms cloning), gRNA sequences and synthetic DNA sequences used in this study. For the primers used to clone left and right homology arms (HAs) using Golden Gate reactions, primer sequences contain BsaI sites in lower case, gRNA binding sites (underlined), and PAMs (bold). For the gRNAs used for landing pad integrations and payload assembly, allele-specific gRNAs are identified as having CAST variants overlapping the PAM, the gRNA seed sequence or both.

This Table is provided as a separate file.

**Table S7. Genomic coordinates.**

Coordinates (mm10) for payloads, genomic locus deletions (marked with a “d”), BACs, enhancer clusters (Enh), and engineered variants. *Igf2*SV, *Igf2* synonymous variants; MI-ICR, methylation-insensitive ICR with CpGs in CTCF sites mutated; and ΔCTCF-ICR, CTCF motif deletions in the ΔCTCF-ICR and locus features. See **Table S5** for payload labels used in figures.

This Table is provided as a separate file.

**Table S8. Summary of payload and mESC capture sequencing libraries.**

Sample ID, a unique sequencing library ID; Payload ID, a descriptor of the relevant landing pad or payload (see **Table S5** for payload labels used in figures); Clone ID, a unique descriptor of mESC clones. For each sample, the number (#) of sequenced, analyzed and duplicate reads is provided. Bait Set is provided for capture samples.

This Table is provided as a separate file.

**Table S9. Payload sequencing coverage QC.**

Summary Table for payload DNA sequencing coverage depth analysis. Length (bp) and normalized (norm) coverage were calculated for regions corresponding to the payload sequence, including different chromosomes for those that apply, as defined in **Methods**. Quality control (QC) calls are included with comments for unexpected values. See **Table S5** for payload labels used in figures.

This Table is provided as a separate file.

**Table S10. Payload variant calling.**

Summary Table for payload variants relative to custom references, related to **Table S2**. Ref, reference sequence; Alt, variant sequence; Qual, quality score; DP, sequencing depth; Feature, the custom reference feature overlapping the variant; RefSeq, variant and surrounding sequence; Comments, annotation and interpretation of variant location and relevance. None of the detected variants are expected to confound the measured activities of integrated payloads. See **Table S5** for payload labels used in Figures.

This Table is provided as a separate file.

**Table S11. mESC clone capture sequencing coverage QC.**

Summary of QC analyses of Capture-seq samples derived from engineered mESC clones, including sequencing coverage, allelic ratios, and integration site analysis (*bamintersect*). See **Methods** for more detail. Samples captured with both *Igf2/H19* and *Sox2* baits (**Table S8**) show borderline low coverage signal at *Sox2* (∼0.65x) due to lower capture efficiency of this last bait, resulting in apparent low payload coverage (marked as FAIL: low coverage [PL]). Clone ID, a unique descriptor of the mESC clone; Sample ID, a unique sequencing library identifier; Payload ID, unique identifier for relevant payload or landing pad. See **Table S5** for payload labels used in figures.

This Table is provided as a separate file.

**Table S12.**
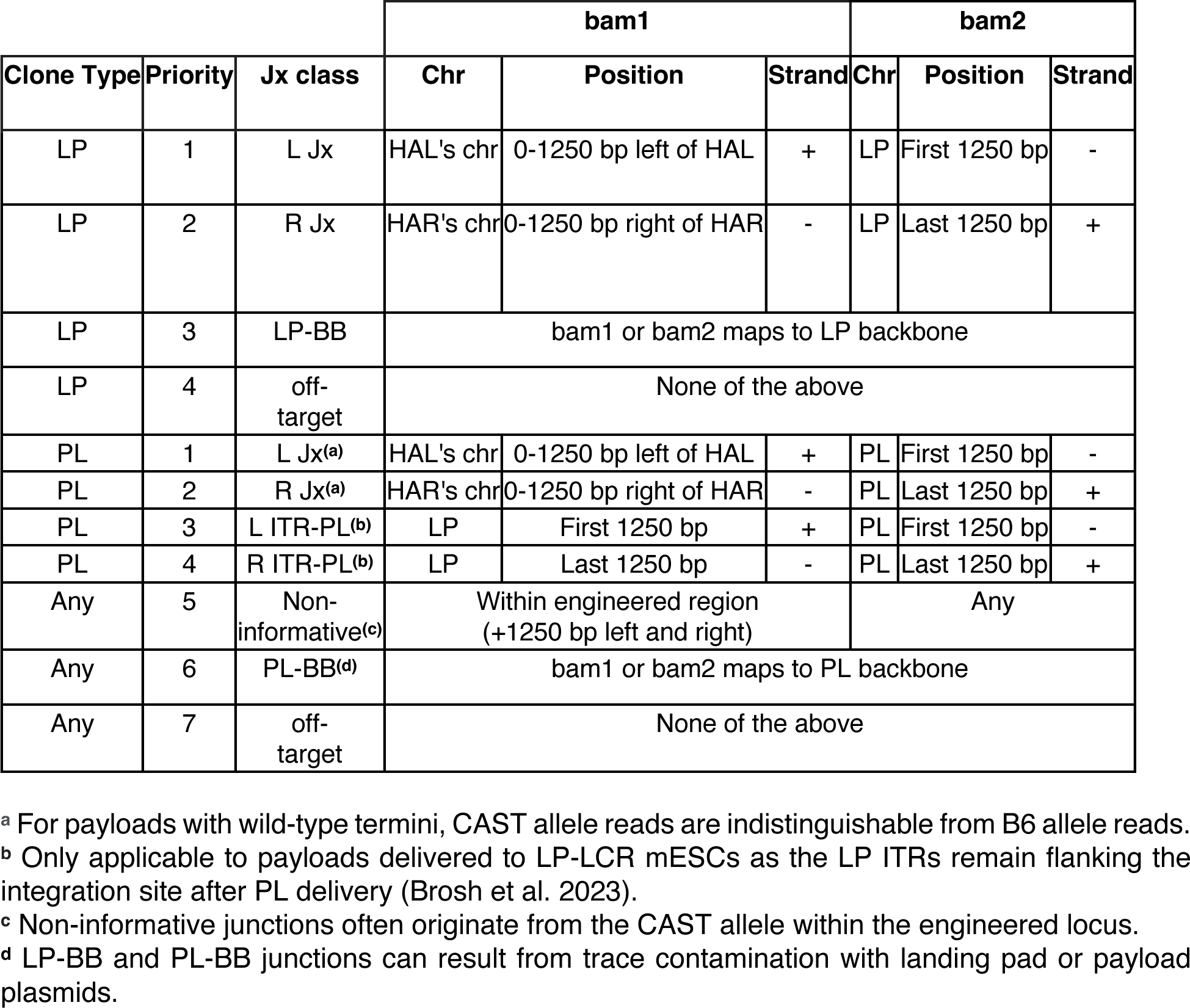
*Bamintersect* junction categories. Hierarchical criteria used to classify *bamintersect* junctions. bam1 and bam2 are the two references to which junction reads were mapped. LP, Landing Pad; PL, payload; BB, backbone; Jx, Junction; Chr, chromosome.

|  |  |  | bam1 |  |  | bam2 |  |  |
| --- | --- | --- | --- | --- | --- | --- | --- | --- |
| Clone Type | Priority | Jx class | Chr | Position | Strand | Chr | Position | Strand |
| LP | 1 | L Jx | HAL's chr | 0-1250 bp left of HAL | + | LP | First 1250 bp | - |
| LP | 2 | R Jx | HAR's chr | 0-1250 bp right of HAR | - | LP | Last 1250 bp | + |
| LP | 3 | LP-BB | bam1 or bam2 maps to LP backbone |  |  |  |  |  |
| LP | 4 | off-target | None of the above |  |  |  |  |  |
| PL | 1 | L Jx<sup>(a)</sup> | HAL's chr | 0-1250 bp left of HAL | + | PL | First 1250 bp | - |
| PL | 2 | R Jx<sup>(a)</sup> | HAR's chr | 0-1250 bp right of HAR | - | PL | Last 1250 bp | + |
| PL | 3 | L ITR-PL<sup>(b)</sup> | LP | First 1250 bp | + | PL | First 1250 bp | - |
| PL | 4 | R ITR-PL<sup>(b)</sup> | LP | Last 1250 bp | - | PL | Last 1250 bp | + |
| Any | 5 | Non-informative<sup>(c)</sup> | Within engineered region (+1250 bp left and right) |  |  | Any |  |  |
| Any | 6 | PL-BB<sup>(d)</sup> | bam1 or bam2 maps to PL backbone |  |  |  |  |  |
| Any | 7 | off-target | None of the above |  |  |  |  |  |

<sup>a</sup> For payloads with wild-type termini, CAST allele reads are indistinguishable from B6 allele reads.
<sup>b</sup> Only applicable to payloads delivered to LP-LCR mESCs as the LP ITRs remain flanking the integration site after PL delivery (Brosh et al. 2023).
<sup>c</sup> Non-informative junctions often originate from the CAST allele within the engineered locus.
<sup>d</sup> LP-BB and PL-BB junctions can result from trace contamination with landing pad or payload plasmids.

**Table S13. mESC clones *bamintersect* results.**

Summary of *bamintersect* integration site analysis for mESC clones capture sequencing samples. Each line describes a junction (Jx) detected by *bamintersect* (see **Methods**). Samples BS17489A and BS17493A missed the left junction due to a known low mappability region overlapping the *Sox2* left homology arm. Samples BS24926A, BS24928A and BS24305A showed low coverage of the right junction, Chrom (chromosome), chromStart, chromEnd, and Strand for bam1 and bam2 refer to the mapped coordinates for the first and second reference genome, respectively. NearestGene indicates the nearest UCSC gene, including distance in bp (negative value indicates an upstream gene) for mm10 or nearest feature for custom references. ReadsPer10M is the number of read pairs supporting the junction per 10 million sequencing reads. Sample ID, a unique sequencing library identifier; Payload ID, LIMS name for relevant payload or landing pad; Clone ID, a unique descriptor of the mESC clone; Genomes indicate the reference genomes compared. Junc_type indicates the junction classification based on criteria detailed in **Table S12**. See **Table S5** for payload labels used in figures.

This Table is provided as a separate file.

## SUPPLEMENTARY DATA

**Data S1. Landing pad and payload sequences.**

Sequences for landing pads and payloads described in this paper, provided in fasta format. For landing pads, sequences include the integrated region (excluding the homology arms). For payloads, sequences include the region from LoxM to LoxP. See **Table S5** for payload labels used in the Figures.

This Table is provided as a separate file.

**Data S2. Raw qRT-PCR Ct values.**

Raw qRT-PCR Ct values for all clones and assays. Payload ID, a unique name for each payload; Label, name as in Figure panels. Location, delivery location in the genome. Additional Deletions, presence of additional deletions engineered in that mESC clone. Label, display name used in Figure panels. Clone ID, a unique identifier for each mESC engineered clone. Cell Type, indicates whether expression was measured in mESC or mesendodermal cells differentiated for 5, 8 or 10 days. Batch indicates the differentiation batch. Assay, targeted gene and allele (engineered or CAST). Cp, threshold cycle.

This Table is provided as a separate file.

